# Splice Factor Polypyrimidine tract-binding protein 1 (Ptbp1) is Required for Immune Priming of the Endothelium in Atherogenic Disturbed Flow Conditions

**DOI:** 10.1101/2021.06.18.449062

**Authors:** Jessica A Hensel, Sarah-Anne E Nicholas, Evan R Jellison, Amy L Kimble, Antoine Menoret, Manabu Ozawa, Annabelle Rodriguez-Oquendo, Anthony T Vella, Patrick A Murphy

## Abstract

NFκB mediated endothelial activation drives leukocyte recruitment and atherosclerosis, in part through upregulation of adhesion molecules Icam1 and Vcam. The endothelium is “primed” for cytokine activation of NFκB by exposure to low and disturbed blood flow (LDF) *in vivo* and by LDF or static conditions in cultured cells. While priming leads to an exaggerated expression of Icam1 and Vcam following cytokine stimulation, the molecular underpinnings are not fully understood. We showed that alternative splicing of genes regulating NFκB signaling occurs during priming, but the functional implications of this are not known. We hypothesize that the regulation of splicing by RNA-binding splice factors is critical for priming. Here, we perform a CRISPR screen in cultured aortic endothelial cells to determine whether splice factors active in the response to LDF participate in endothelial cell priming. Using Icam1 and Vcam induction by TNFα stimulation as a marker of priming, we identify polypyrimidine tract binding protein (Ptbp1) as a required splice factor. Ptbp1 expression is increased and its motifs are enriched nearby alternatively spliced exons in endothelial cells exposed to LDF *in vivo* in a platelet dependent manner, indicating its induction by early innate immune cell recruitment. At a mechanistic level, deletion of Ptbp1 inhibited NFκB nuclear translocation and transcriptional activation. These changes coincided with altered splicing of key components of the NFκB signaling pathway that were similarly altered in the LDF response. However, these splicing and transcriptional changes could be restored by expression of human PTBP1 cDNA in Ptbp1 deleted cells. *In vivo,* endothelial specific deletion of Ptbp1 reduced myeloid cell infiltration at regions of LDF in atherosclerotic mice. In human coronary arteries, PTBP1 expression correlates with expression of TNF pathway genes and amount of plaque. Together, our data suggest that Ptbp1, which is activated in the endothelium by innate immune cell recruitment in regions of LDF, is required for priming of the endothelium for subsequent NFκB activation and myeloid cell recruitment in vascular inflammation.

**Graphical Abstract:** Plaque forms in low and disturbed flow regions of the vasculature, where endothelial cells are “primed” to respond to cytokines (e.g. TNFα) with elevated levels of cell adhesion molecules via the NFκB signaling pathway. We show that the splice factor Ptbp1 (purple) mediates priming. Ptbp1 is induced in endothelial cells by platelet recruitment, promoting priming and subsequent myeloid cell infiltration into plaque. Mechanistically, Ptbp1 regulates splicing of genes involved in the NFκB signaling pathway and is required for efficient nuclear translocation of NFκB in endothelial cells. This provides new insight into the molecular mechanisms underlying an endothelial priming process that reinforces vascular inflammatory responses.

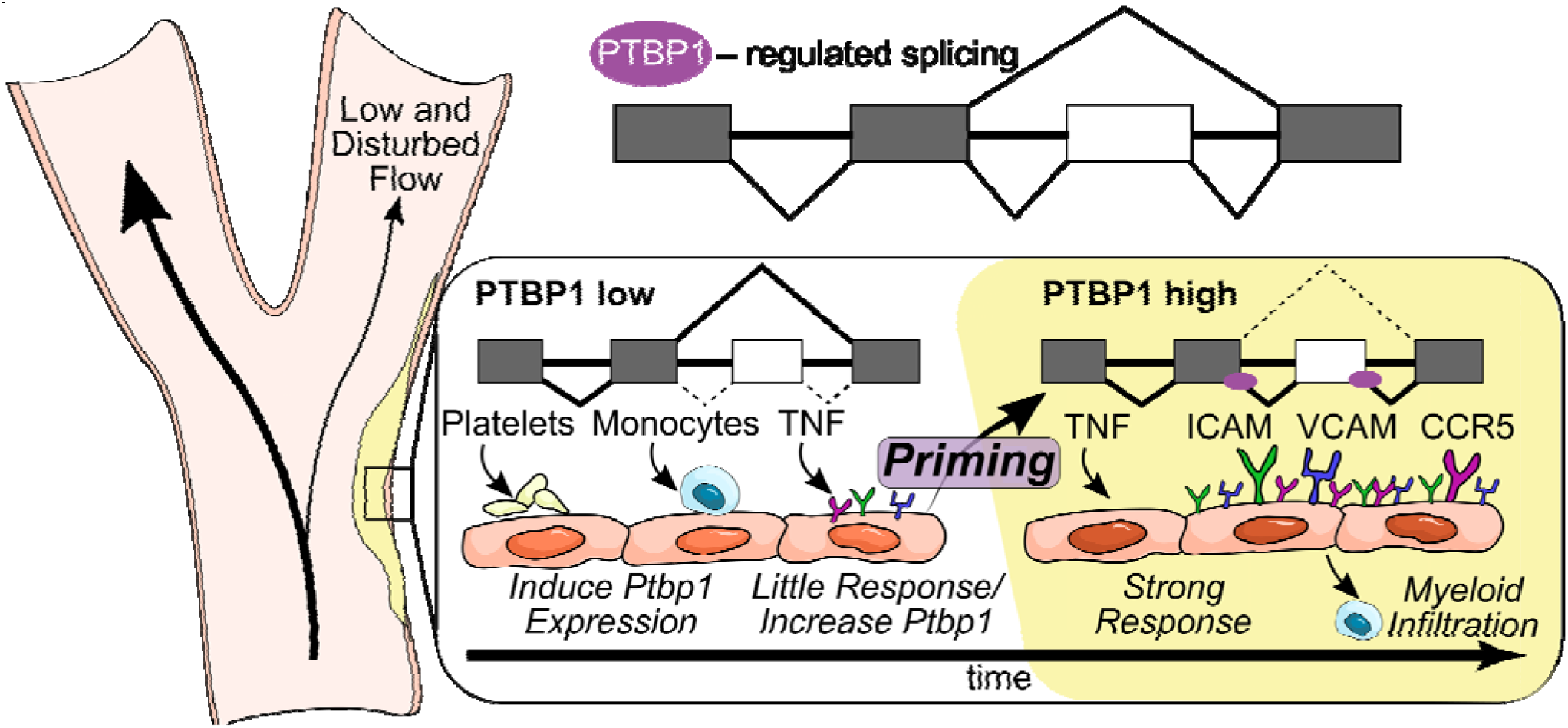

## Introduction

Atherosclerosis is a disease characterized by chronic sterile inflammation, driven by leukocyte recruitment to regions of low and disturbed flow (LDF) in arteries. In these regions, endothelial expression of leukocyte adhesion molecules, such as Icam1, Vcam, and P-,E- and L-selectins, and cytokines, such as CCL2 (MCP1) and CCL5 (RANTES), mediate leukocyte recruitment ^1,2^. Complete inhibition of these molecules, or the NFκB signaling pathway suppresses leukocyte recruitment to the endothelium, and reduces both acute and chronic inflammation in atherosclerotic plaque ^3^. The expression of these molecules is tightly regulated in the endothelium, primarily by NFκB signaling. Activity of this pathway and expression of downstream leukocyte recruitment factors is low under quiescent conditions, but increased through both transcriptional and post-transcriptional mechanisms in response to circulating inflammatory cytokines. Not all areas of the vasculature are similarly affected however, as it is only regions exposed to LDF that exhibit elevated levels of adhesion molecules in response to NFκB agonists ^4^.

While low and disturbed flow alone is insufficient to induce high levels of adhesion molecule expression, it does prime the endothelium for subsequent activation by cytokines. Endothelial cell priming in disturbed flow regions *in vivo* is demonstrated by response to systemic NFκB agonist lipopolysaccharide (LPS) with increased Vcam and E-selectin expression, while cells in regions with laminar flow do not ^4^. *In vitro*, endothelial cells cultured under LDF or static conditions respond to NFκB agonist Il1β by increasing surface Vcam expression, while cells cultured under laminar flow conditions do not ^5^. Both chronic and acute changes in the endothelium contribute the priming of NFκB responses. In chronic conditions of LDF, subendothelial deposition of fibronectin changes the repertoire of integrin binding and increases NFκB responses ^6–8^. However, the mechanisms responsible for priming following acute changes in flow are poorly understood and may contribute to atherosclerosis and other inflammatory responses.

Using an acute partial carotid ligation model of LDF, we observed alterations in post-transcriptional regulation by RNA splicing in primed endothelial cells ^9^. Among the affected transcripts were multiple regulators of the NFκB signaling pathway, including canonical pathway genes such as Iκbκγ (Nemo) and other known regulators, such as fibronectin ^9^. Alternative splicing, which is coordinated by the spliceosome but directed by hundreds of RNA-binding proteins in the cell, allows a single gene and pre-mRNA to encode multiple unique mRNA molecules and protein products. Alterations in the levels and activities of RNA-binding splice factors can allow a cell to rapidly modify the composition of signaling pathways. For example, increased inclusion of alternative exons EIIIA and EIIIB in fibronectin is linked to alterations in extracellular matrix composition, integrin binding, and increased NFκB signaling ^7,10^. Thus, we hypothesize that alternative splicing mediates acute endothelial priming during the arterial response to LDF. Here, we test this hypothesis by determining whether splice factors of the arterial endothelium activated in response to LDF are important in endothelial cell priming, identifying Ptbp1 as a potent mediator of priming and subsequent myeloid cell recruitment and atherogenesis.

## Results

### A CRISPR-KO screen of splice factors modulating alternative splicing in endothelial cells exposed to low and disturbed flow reveals a requirement for Polypyrimidine tract-binding protein 1 (Ptbp1) in endothelial cell priming

To identify splice factors activated in endothelial cells in the earliest stages of priming, we examined splicing patterns in endothelial cells experimentally exposed to LDF for 48hrs through a partial carotid ligation (Figure 1A)^9^. In our earlier work, we had observed that many of the splicing changes in the endothelium were dependent on platelet recruitment. Platelets are an essential component of endothelial cell priming as regions of LDF do not recruit myeloid cells in the absence of platelets ^11–14^. Additionally, activated platelets are sufficient to increase endothelial Icam and Vcam in static *in vitro* culture conditions that mimic the effects of LDF ^15^. Thus, we focused on splicing changes induced in the endothelium upon platelet recruitment. Of the several thousand splicing changes we detected in the intima in priming (Figure 1B), we found that most were reverted by the depletion of platelets by anti-GPIBα (Figure 1C, D and E, and SI Figure 1). To determine which splice factors might be regulating platelet induced splicing changes under LDF, we assembled a list, based on (i) differential splice factor expression, (ii) enrichment of motifs nearby regulated core exons (CE), or (iii) splicing of the factor itself, which is often an indication of splice factor activity and autoregulation ^16^. Based on these criteria, we deemed 57 splice factors likely to be important in the regulation of splicing changes in the endothelium under these conditions (Figure 2A).

**Figure 1.**
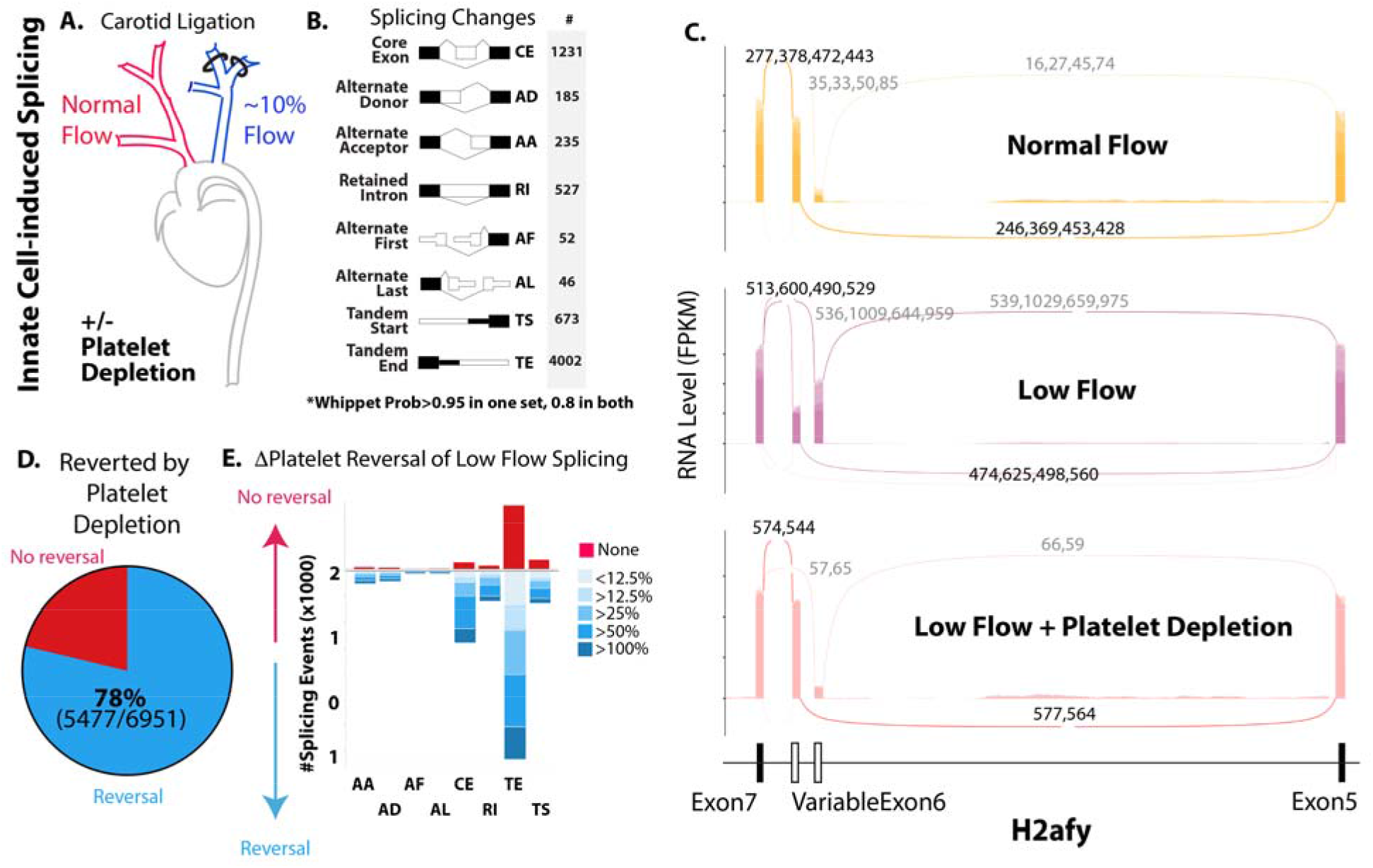
Depletion of platelets reverts much of the splicing response induced in the arterial endothelium by exposure to low and disturbed flow (LDF). (A) Schematic of approach for testing RNA splicing responses induced by acute exposure to low and disturbed flow (LDF), (*Murphy et al*. eLife *2018*). Ligation of distal branches of the left carotid artery (blue) leads to a 10% reduction in blood flow. Platelets were depleted by anti-GPIBα treatment at the time of the surgery. (B) Change in splicing analyzed by Whippet. Changes shown represent those with a probability of > 0.95 in one low flow comparison (N=2 v N=2 pools, each pool containing 2-3 arteries), and a probability of at least 0.8 in two separate experiments (N=2 v N=2 pools each). (C) Examples of splicing events in H2afy RNA induced by LDF and reversed by the depletion of platelets. Sashimi plot shows the read density (y-axis) and number of splice junction spanning reads. Low flow and high flow each N=4 (each a pool of multiple vessels), low flow + platelet depletion N=2 (each a pool of multiple vessels). The number splice junction reads in each biological replicate are separated by commas. (D&E) Summary of the degree of reversal (or not) with platelet depletion of all splicing events defined in panel B induced by LDF (D) and then disaggregated by type of splicing events, with a scale to indicate the degree of reversal (E).

**Figure 2.**
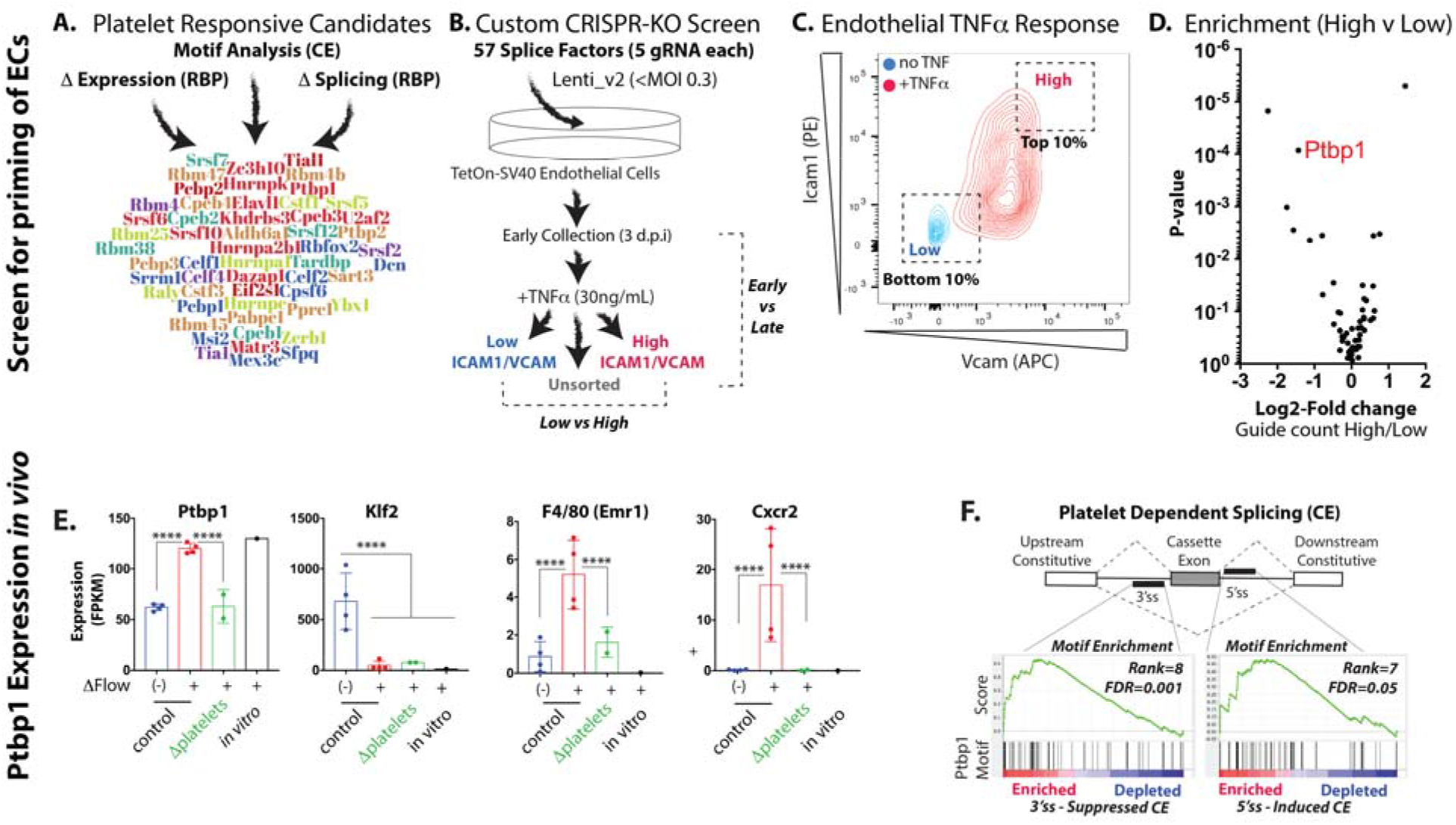
CRISPR-KO screen to examine splice factor involvement in endothelial priming. (A) Summary of the development of a list of splice factors that may play an important role in endothelial cell priming by exposure to LDF. CE=core exon. RBP=RNA-binding protein. (B) *In vitro* screen to determine which of the candidate splice factors regulate endothelial priming - defined here by upregulation of Icam1 & Vcam following exposure to TNFα. d.p.i.=days post-infection with CRISPR library. (C) Example flow cytometry plot, showing Icam1 and Vcam protein expression in the CRISPR KO pool or aortic endothelial cells before and after treatment with TNFα. (D) MAGeCK analysis of differentially detected guide RNA in cells with high Icam1 & Vcam response versus low response, grouped by gene (5 gRNA per gene). Enrichment in guide count (x-axis) and P-value (Y-axis) are shown. (E) Changes in the levels transcripts from the indicated genes in the carotid intima of vessels exposed to LDF (ΔFlow+), or from normal flow contralateral control (ΔFlow(-)), or from cultured cells under static *in vitro* conditions (*in vitro*) by RNA-seq analysis. Each point represents a pool of 2-3 arteries. FPKM=Fragments Per Kilobase of transcript per Million mapped reads. (F) Kolmogorov-Smirnov based enrichment of *in vitro* defined Ptbp1 6-mer motifs nearby LDF and platelet regulated core exons, relative to skipped exons which were not altered by the loss of platelets under LDF conditions. Rank displayed is relative to all other splice factor motifs searched. (E) ****Padj<0.0001, by DESeq2.

To assess contributions of these splice factors to endothelial cell priming, we established an *in vitro* CRISPR-KO screening approach which took advantage of our previous discovery that splicing patterns in cultured aortic endothelial cells resemble endothelial cells primed by LDF *in vivo* (Figure 2A & SI Figure 2&3). In our screen, we treated aortic endothelial cells with TNFα, and examined induction of Icam1 and Vcam. Guide RNA (gRNA) targeting genes important in priming should be found more often in cells with low Icam1 & Vcam responses than high responses, indicating that deletion of the targeted splice factors suppressed the TNFα-response. Indeed, we could identify cells in our CRISPR pool that did not respond strongly to TNFα (Figure 2C). Importantly, we observed that this “low responder” phenotype was retained upon restimulation, indicating their stable alteration by CRISPR editing (SI Figure 4). Using this approach, and subsequent identification of enriched gRNA in low and high responders by 1) PCR amplification of lentiviral insertions, 2) high-throughput sequencing and 3) statistical analysis, we found that Ptbp1 gRNA was consistently enriched in cells with a reduced Icam1 & Vcam response to TNFα across multiple screens (Figure 2D, and SI Table 1). We also observed that gRNA to Ptbp1 was enriched in low responders in our iterative sorting experiment (low responders plated and sorted for low response a second time, (SI Table 1).

To determine whether impaired Icam1 & Vcam induction in cells containing Ptbp1 gRNA might be a result of generally impaired cell function, we examined presence of Ptbp1 gRNA at early and late timepoints in our pool of cells. Losses could represent reduced cell viability or proliferation, while gains could reveal growth-suppressive splice factors as deletion of these genes would allow these cells to expand within the pool over time. We found no significant loss or enrichment of Ptbp1-targeting gRNA, while we did observe reduced representation of other gRNA and increased representation of gRNA for Hnrnpa1, a splice factor that has been shown to inhibit cell proliferation in other contexts (SI Figure 5 and SI Table 2)^17^.

We then reexamined the expression and motif analysis (Figure 2A) that had landed Ptbp1 on our short list of splice factors to be targeted, and observed that Ptbp1 expression is induced, in a platelet dependent manner, by exposure of the carotid endothelium to LDF (Figure 2E). There is a similar induction in endothelial cells in static conditions and in the presence of serum — as we would expect under these priming *in vitro* conditions (Figure 2E). Expression of Klf2, a transcription factor reduced in expression under LDF, was not affected by platelet depletion, while myeloid cell recruitment (indicated by macrophage marker F4/80 and neutrophil marker Cxcr2) was strongly impaired. These data support the hypothesis that platelets are not required for the initial low flow response of the endothelium (e.g. Klf2), but are required for full immune-activation. Finally, we found significant enrichment of Ptbp1 motifs at the 3’splice site side of core exons (CE) that were included less frequently under LDF conditions in a platelet dependent manner, consistent with the known role of increased Ptbp1 in suppressing exon inclusion from the 3’splice site (Figure 2F)^18,19^. We also found a weaker association of Ptbp1 motifs with the 5’ splice site side of CE that were included more frequently under LDF in a platelet dependent manner (Figure 2F).

Thus, informatics analysis and a targeted CRISPR-KO screen suggest that endothelial Ptbp1 expression, which is induced in a platelet dependent manner under LDF, contributes to the priming of endothelial cells.

### Endothelial Ptbp1 is required for NFκB-mediated transcriptional activation

The dampening effect of Ptbp1 deletion on TNFα-induced expression of Icam1 and Vcam on the surface of endothelial cells could be mediated by several mechanisms, including a block in NFκB transcriptional activity or inhibition of Icam1 & Vcam translation or translocation to the plasma membrane. We had previously observed changes in splicing in key components of the NFκB signaling pathway in the response to LDF (e.g., IκBκγ, or Nemo) ^9^, so we focused on NFκB transcriptional activation.

To examine NFκB transcriptional activity, we modified a lentiviral NFκB reporter ^20^, introducing eGFP as the reporter gene downstream of the NFκB binding motif (Figure 3A). Using this reporter, we again screened a CRISPR KO pool of aortic endothelial cells, with our targeted group of splice factors. Ptbp1 gRNA was enriched in cells with reduced NFκB activity to TNFα as determined by decreased GFP fluorescence (Figure 3B), suggesting that deletion of Ptbp1 impairs TNFα-mediated NFκB transcriptional responses.

**Figure 3.**
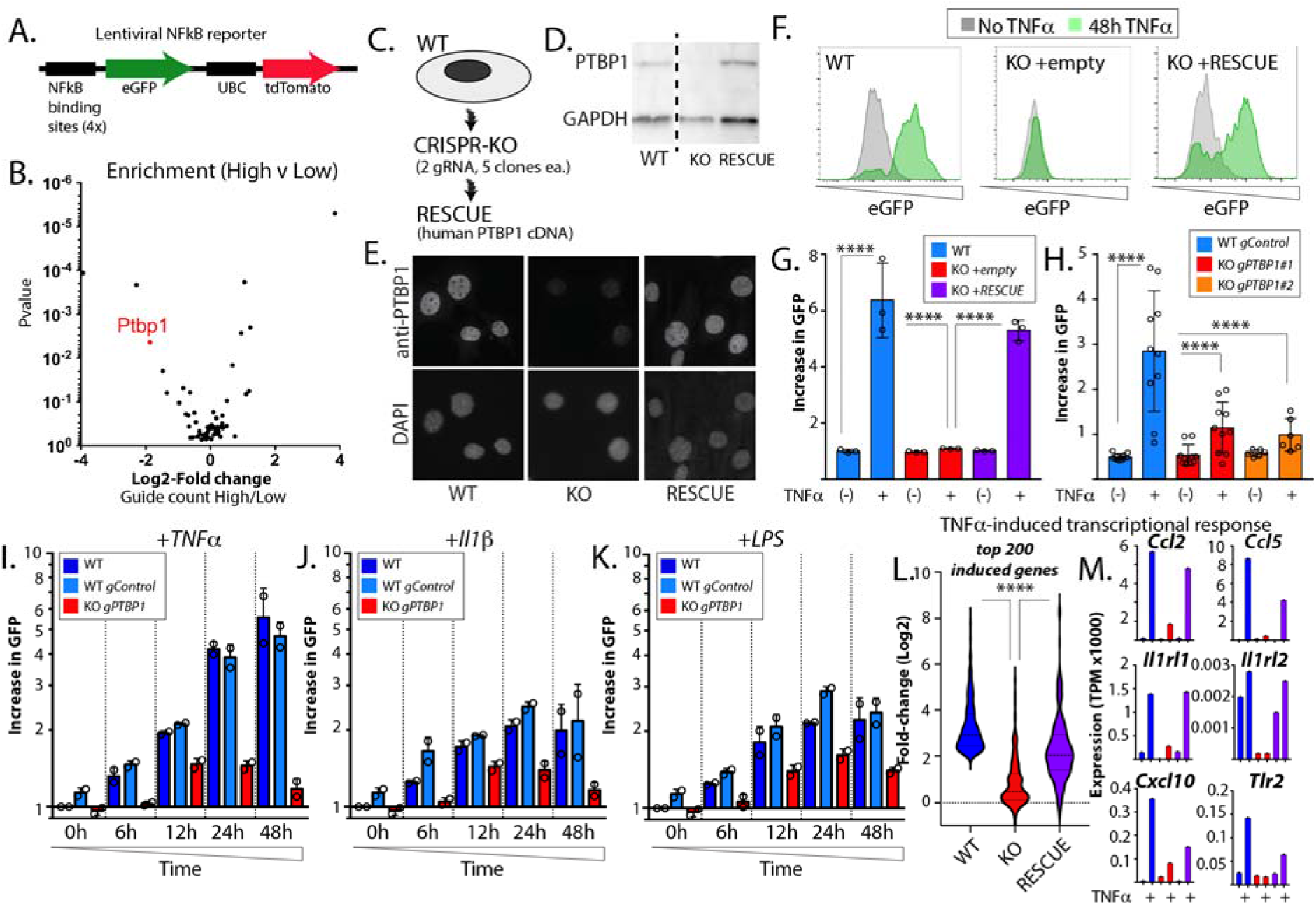
Endothelial Ptbp1 is required for NFκB-mediated transcriptional responses. (A) Engineered lentiviral NFκB reporter, used to assess NFκB transcriptional activity. UBC=ubiquitin C promoter, used to drive constitutive tdTomato expression - a control for Lentiviral expression in the cells. (B) MAGeCK analysis of results of CRISPR KO pool, using NFκB reporter to examine response to TNFα. (C) Schematic showing approach to determine the effect of endothelial Ptbp1 on endothelial “priming.” (D) Western blot and (E) immunostaining of cells, showing antibody staining of PTBP1 (D&E) and GAPDH (D) in a representative CRISPR-KO single-cell clone and this clone with re-expression of human PTBP1 cDNA. (F) Single-cell clone of Ptbp1 CRISPR KO, showing NFκB response by reporter, and restoration of TNFα response by human PTBP1 cDNA rescue construct. (G&H) Quantitation of mean increase in eGFP expression in each cell line after TNFα treatment. (G) N=3 biological replicates. (H) N=2 biological replicates of N=5 single cell clones (gControl) or Ptbp1 targeting guide #1 (gPTBP1#1), N=2 biological replicates of N=3 single cell clones gPTBP1#2. (I-K) Quantitation of mean increase in eGFP over time for cells with NFκB reporter only, cells with reporter + control guide (gControl), or + Ptbp1 targeting guide (gPTBP1). Stimulation at 0 hour with (I) 30 ng/mL TNFα, (J) 30 ng/mL Il1β, or (K) 30 ng/mL LPS. (L) Analysis of the top 200 TNFα-induced transcripts in wild-type cells, Ptbp1 KO cells, and rescue cells, as determined by RNA-sequencing before and after 24 hours of 30 ng/mL TNFα. (M) Examples of transcripts from L, showing expression levels. (G,H,&L) ****P<0.001, Kruskal-Wallis with Dunn’s multiple comparison test.

CRISPR activity in pooled approaches leads to varying levels of deletion within the population, so to assess the effects of a verified Ptbp1 deletion, we generated independent CRISPR-KO endothelial cell clones, using two different Ptbp1 gRNA, as well as a set of non-targeting gRNA single cell clones (Figure 3C). To further confirm specificity, we rescued Ptbp1 expression in these Ptbp1 KO clonal lines by introducing a human PTBP1 cDNA construct (Figure 3C and data not shown). This is 97% identical to murine Ptbp1 at the amino acid level, but prevents CRISPR targeting of the construct due to the divergent nucleic sequence. We demonstrated Ptbp1 loss in the KO lines, and found that the human PTBP1 rescue restored Ptbp1 expression (Figure 3D) and nuclear localization (Figure 3E).

Using these Ptbp1 KO cell lines, we examined our NFκB transcriptional reporter and found that the NFκB response to TNFα was blocked by the loss of Ptbp1, and restored upon human PTBP1 rescue (Figure 3F with quantification in 3G). Multiple single-cell clones from different gRNA exhibited the same response, further verifying these results (Figure 3H). Notably, as we had previously observed in our screens, loss of Ptbp1 had little effect on endothelial cell survival or proliferation, as we found no significant differences in proliferation among these different cell lines (SI Figure 5B).

The NFκB signaling pathway is a common downstream mediator of many upstream inputs, including TNFα, Il1β, and LPS, each of which signal through different receptors at the plasma membrane. To understand whether Ptbp1-mediated NFκB inhibition was specific to TNFα, or whether it may act on the core pathway downstream of multiple inputs, we examined eGFP responses at multiple timepoints in CRISPR-KO cells and both wild-type or gRNA control cells. We found a similar inhibition of all signaling responses, beginning as early as 6 hours (Figure 3I-K), suggesting that loss of Ptbp1 inhibits core NFκB signaling.

As our work had focused on a single synthetic NFκB responsive element, we wondered if endogenous NFκB-mediated transcription was similarly affected. To examine this, we performed RNA-sequencing analysis on wildtype cells, our Ptbp1 CRISPR-KO cell line, and human PTBP1 rescue of these Ptbp1 CRISPR-KO cells, before and after treatment with TNFα. As expected, TNFα treatment led to large changes in transcription of canonical NFκB target genes. Analysis of the top 200 induced transcripts found in wild-type cells showed a nearly complete block in induction by TNFα in the Ptbp1 CRISPR-KO cells, and that this could be rescued, although not completely, in the human PTBP1 rescue cells (Figure 3L&M). The partial rescue could reflect differences in human and mouse Ptbp1, or that we restored only a single isoform of Ptbp1 - as we noted a second Ptbp1 band on Western blots that was not restored with human cDNA expression.

Thus, Ptbp1 is critical for NFκB signaling responses in endothelial cells downstream of various inputs. Expression of Ptbp1 primes endothelial cell to respond to TNFα, Il1β, and LPS, licensing them to express a wide range of genes involved in the regulation of inflammation and immunity.

### Ptbp1 coordinates splicing in endothelial activation that modulates nuclear translocation of NFκB

We reasoned that if Ptbp1 affects the core NFκB signaling pathway, it may alter expression or splicing of key components or modifiers of the pathway. To test this, we assessed levels of core NFκB signaling genes, using RNA-sequencing data from the different cell lines generated (Figure 4A). With the exception of Ripk1 and Nfkbia, we found very little change in any of these genes at the whole transcript level, despite strong effects on canonical NFκB-induced genes. We then examined changes in RNA splicing. Consistent with the splice factor activity of Ptbp1, we observed many changes in alternative splicing, including some in coding regions of core NFκB pathway genes (e.g., IκBκγ/NEMO & Ubc), and key regulators of the NFκB signaling pathway (e.g., Pidd1, Stat3, Irak4, and Pdlim7) (Figure 4B&C and SI Table 4). In total, there were 110 splicing changes in the Kegg NFκB pathway genes, 54 with a splicing difference of >10% and 38 with a splicing difference >20% (SI Table 4). In these datasets, splicing levels at a particular junction are indicated by the percent spliced in (Psi), where a Psi of 0.1=10% spliced and 0.9=90% spliced in. Notably, large changes in RNA splicing often occurred in the absence of changes in total RNA expression, as was the case for IκBκγ (Figure 4B&D). For other genes, like Pdlim7, changes in splicing were correlated, though generally larger than the changes in transcript levels.

**Figure 4.**
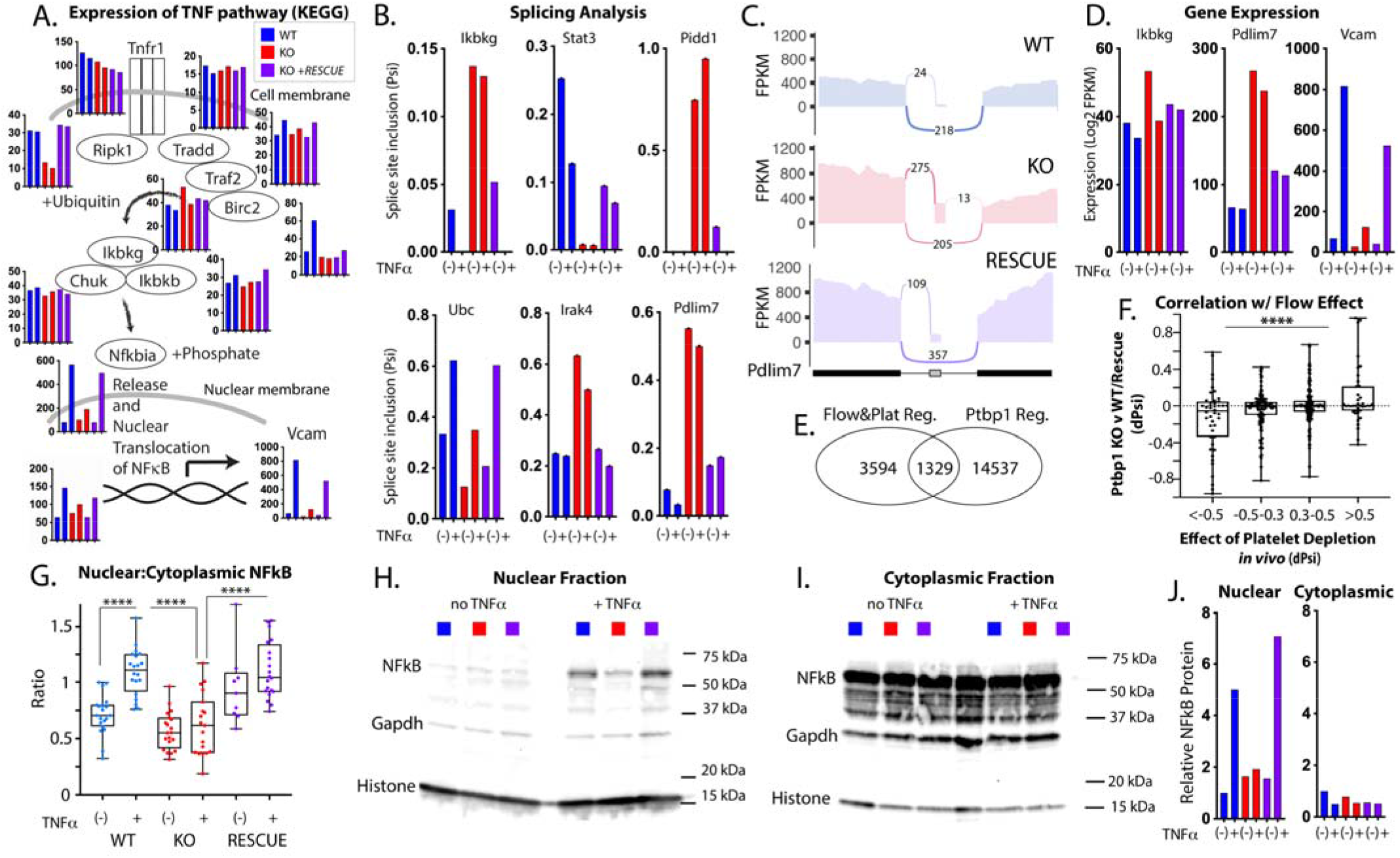
Ptbp1 is required for the nuclear translocation of NFκB and modulates splicing of key components and regulators of the NFκB pathway. (A) Analysis of gene expression (Transcripts per Million, TpM) in the indicated cell lines for components of the canonical NFκB signaling pathway, by RNA-sequencing before or after 24h treatment with 30ng/mLTNFα. Organization of the bars is the same as (B &D). (B) Examples of Ptbp1 regulated splicing events in the canonical NFκB signaling pathway, or in genes regulating the NFκB signaling pathway, as analyzed by Whippet in cultured cells with or without Ptbp1. Splice site inclusion (Psi) indicates the fraction included. (C) Sashimi plots, showing examples of differentially spliced transcripts of Pdlim7. (D) Select genes in the NFκB pathway, or known to regulate NFκB signaling, from this same data set. (E) Venn diagram showing the overlap between the splicing events regulated by platelets in the *in vivo* endothelium and those regulated by Ptbp1 *in vitro* (present in wildtype or rescue cells, but not KO); at least 5 reads were required in all data sets (Probability of 0.8 in three different *in vivo* data sets for platelet regulation, and 0.9 in two different *in vitro* comparisons - all under TNFα treatment, wildtype cells vs Ptbp1 KO, and Ptbp1 KO cells vs Ptbp1 KO cells + human PTBP1 rescue). (F) Analysis of all platelet regulated splicing events, comparing the change in splicing inclusion (Psi) with platelet depletion *in vivo* to the change observed with Ptbp1 deletion *in vitro*. (G) Quantitation of nuclear translocation of NFκB by immunofluorescence, indicated as the ratio between nuclear and cytoplasmic staining in cultured cells in the indicated cell lines, before and after 24h treatment with 30ng/mL TNFα. (H&I) Western blots from nuclear (H) and cytoplasmic (I) enriched fractions from aortic endothelial cells from the cell lines indicated in the key, with or without 24h treatment with 30ng/mL TNFα. (J) Quantification of signal, relative to control (Histone in nuclear and Gapdh in cytoplasmic), and normalized to wildtype cells. (F&G) ****P<0.001, Kruskal-Wallis with Dunn’s multiple comparison test.

To determine whether Ptbp1 affected splicing of genes in the platelet-mediated endothelial activation we previously observed (Figure 1), we examined the overlap between platelet regulated transcripts *in vitro* and this *in vivo* dataset, limiting to splicing events with at least 5 reads of support in both sets. We found evidence for Ptbp1 regulation of ~25% of all the platelet-regulated splicing events (Figure 4E). Analysis of these splicing events revealed that those with reduced inclusion upon platelet depletion also showed reduced inclusion upon Ptbp1 depletion (Figure 4F), and vice versa, consistent with our hypothesis that Ptbp1 activity is increased in endothelial cells under LDF in a platelet dependent manner.

To determine whether Ptbp1 loss affected localization or activity of NFκB components, we examined the expression of key proteins in the pathway, including IKKα, IKKβ, IκBκγ(NEMO), IKβα and its phosphorylation, and nuclear translocation of NFκB. Although levels of IKKα, IKKβ, IκBκγ, and IKβα were similar (SI Figure 6), we observed reduced TNFα-mediated nuclear translocation of NFκB in Ptbp1 KO cells, which was rescued by restoration of Ptbp1 (Figure 4G-J).

Thus, loss of Ptbp1 altered splicing in several genes in the core NFκB signaling pathway (e.g. IκBκγ), or in important modifier pathways (e.g., Pdlim7), and lead to a loss of NFκB nuclear translocation following TNFα treatment.

### Endothelial Ptbp1 deletion reduces myeloid cells in atherosclerotic plaque

To test the *in vivo* requirement for Ptbp1 in endothelial cell priming in the context of atherosclerosis, we generated a conditional endothelial specific Ptbp1 mouse (*Cdh5(PAC)CreERT2; Ptbp1^ff^* or EC-KO), by intercrossing mice with these alleles ^21,22^. We induced Ptbp1 excision in 6 to 7 week-old mice by tamoxifen treatment. We induced hypercholesteremia by treating Ptbp1 EC-KO mice or their littermate controls with AAV-PCSK9 and a high fat diet^23^. After 3 months of this diet, we collected aortic arches, innominate arteries and blood samples for immune cell analysis and plaque composition. The inner curvature of the aortic arch and the innominate arteries are regions exposed to LDF, where endothelial cells are primed and plaque develops ^4^. We found that, while both Ptbp1 EC-KO mice and their littermate controls exhibited a robust increase in cholesterol levels (~10-fold standard chow diet mice; Figure 5B) and developed atherosclerotic plaque in the aortic arch, that plaque was visibly reduced in the arches of EC-KO mice (Figure 5C&D). Consistent with an impaired inflammatory priming of the endothelium, there were also fewer CD11b+ cells found in these plaques (Figure 5E), and the ratio of CD11b+ cells to CD45+ cells was reduced in the plaque (Figure 5F). We observed a similar trend in CD11b+Gr1+ cells (Figure 5G), although the proportion of CD11b+ cells that were also Gr1+ was very low (~10%). This did not appear to result from a systemic reduction in the number of circulating CD11b+ cells, either in total, or as a percentage of CD45+ cells, as these were not reduced in Ptbp1 EC-KO mice (Figure 5H-J).

**Figure 5.**
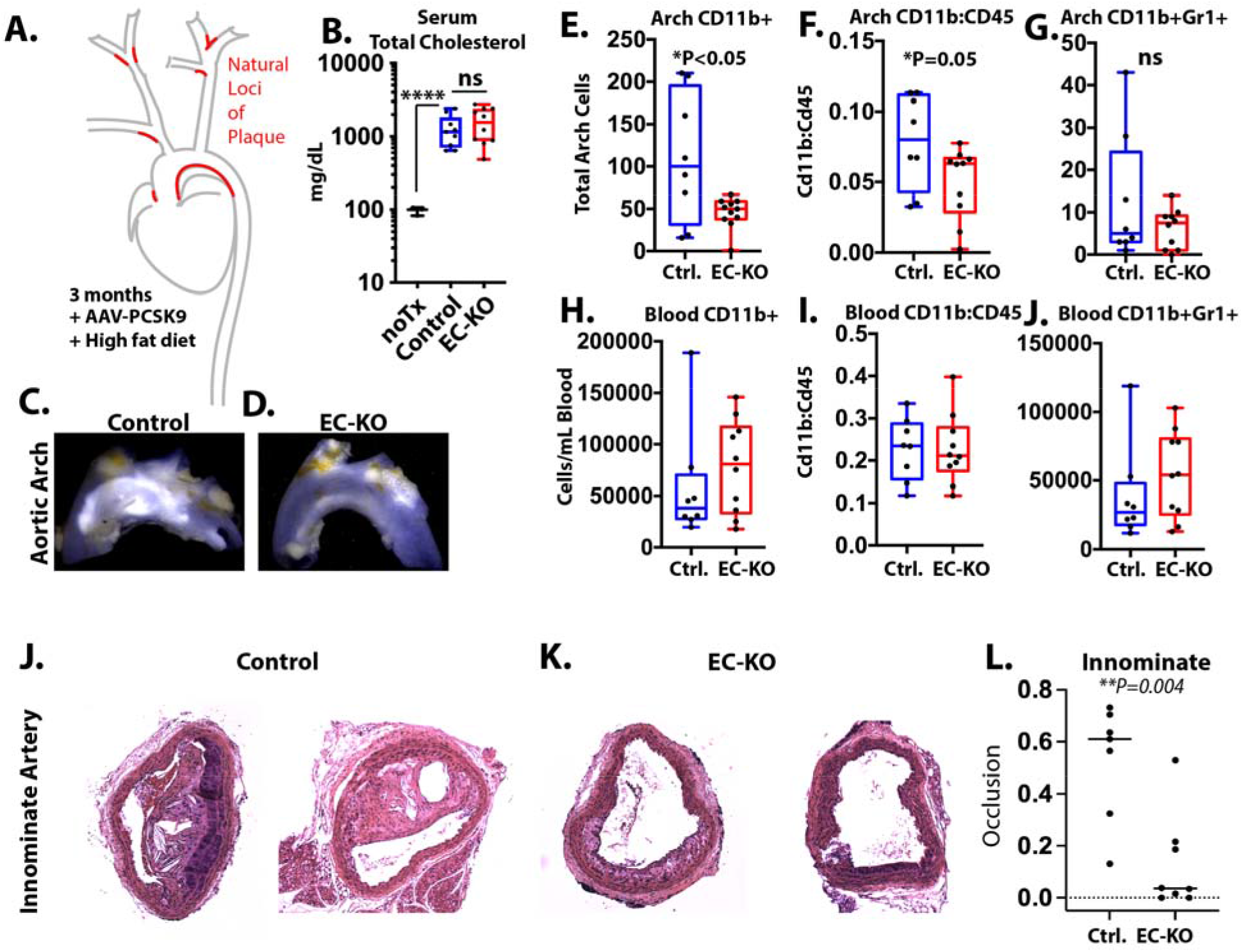
Hypercholesteremic Ptbp1 EC-KO mice exhibit impaired recruitment of myeloid cells to the aortic arch. (A) Schematic of model, showing treatment protocol with AAV-PCSK9 + high-fat diet (HFD) and LDF sites of plaque deposition (red). (B) Serum cholesterol levels in untreated mice (noTx) and the Ptbp1 EC-KO mice and their littermate controls (EC-KO and Control), 3 months after treatment with AAV-PCSK9 and HFD. (C&D) Representative gross images of aortic arch from (D) EC-KO mice or (C) littermate controls when isolated 3 months after AAV-PCSK9 and HFD. White regions in these arches are plaque. (E-J) Analysis of myeloid cells in the aortic arch and the blood of these mice, showing total cells per arch or mL of blood, the ratio of CD11b+ to CD45+ cells, and the total number of CD11b+Gr1+ cells. (J-L) H&E images of representative innominate arteries of the indicated genotypes just before the brachiocephalic branch point, and (L) Quantitation of the degrees of vessel occlusion (1=100%, 0=0%). P-value from Mann-Whitney test.

Therefore, Ptbp1 deletion from the endothelium impairs myeloid cell accumulation in atherosclerotic plaque in regions of endothelial cell priming, consistent with the reduced levels of priming we observed in cultured cells.

### Ptbp1 expression positively associates with plaque burden and TNF/NFκB signaling pathway in human arteries

We reasoned, if Ptbp1 is required for efficient TNFα/NFκB signaling responses and myeloid cell recruitment to plaque, that we might find these correlations in human tissues. To test this, we examined the GTEx database, which contains RNA transcript analysis and pathology reports from hundreds of donor arteries (Figure 6A). While this database is focused on non-diseased tissues, nearly all arteries in the donor group collected include some degree of plaque formation, reflecting the ubiquitous nature of this process in human arteries with age. Nevertheless, a small subset of arteries was deemed free, or mostly free of plaque by pathology report (Figure 6B), while others had increasing levels of plaque (Figure 6C). As expected, expression of TNFα and Il1β, cytokines central to plaque development and immune cell composition, were enriched in arteries with plaque versus those without (Figure 6D&E). Ptbp1 and the TNF receptor were also increased in arteries with plaque versus those without (Figure 6F&G). Notably, there is a wide range of expression of both cytokines and their receptors in plaque, reflecting widely varied immune milieus within plaque (Figure 6D-G). This variation is important, as immune cell composition in plaque is strongly correlated with myocardial infarction and stroke. Therefore, we asked what pathways are most associated with Ptbp1 in coronary arteries by examining gene-gene correlations with Ptbp1 expression (e.g. TNFRSF1A, Figure 6H). Doing this for all genes in specific arteries (e.g. coronary, tibial, aorta) revealed a strong conservation of gene-gene interactions between arteries (Figure 6I), but that genes correlated with Ptbp1 in arteries did not have the same correlation in whole blood (Figure 6J). We then asked what gene groups are most often positively correlated with Ptbp1 expression, and found a strong signature for Myc (which can induce Ptbp1 expression, and is also regulated by Ptbp1 ^24–26^), but also Interferon gamma signaling, and TNF signaling via NFκB (Figure 6K&L and SI Table).

**Figure 6.**
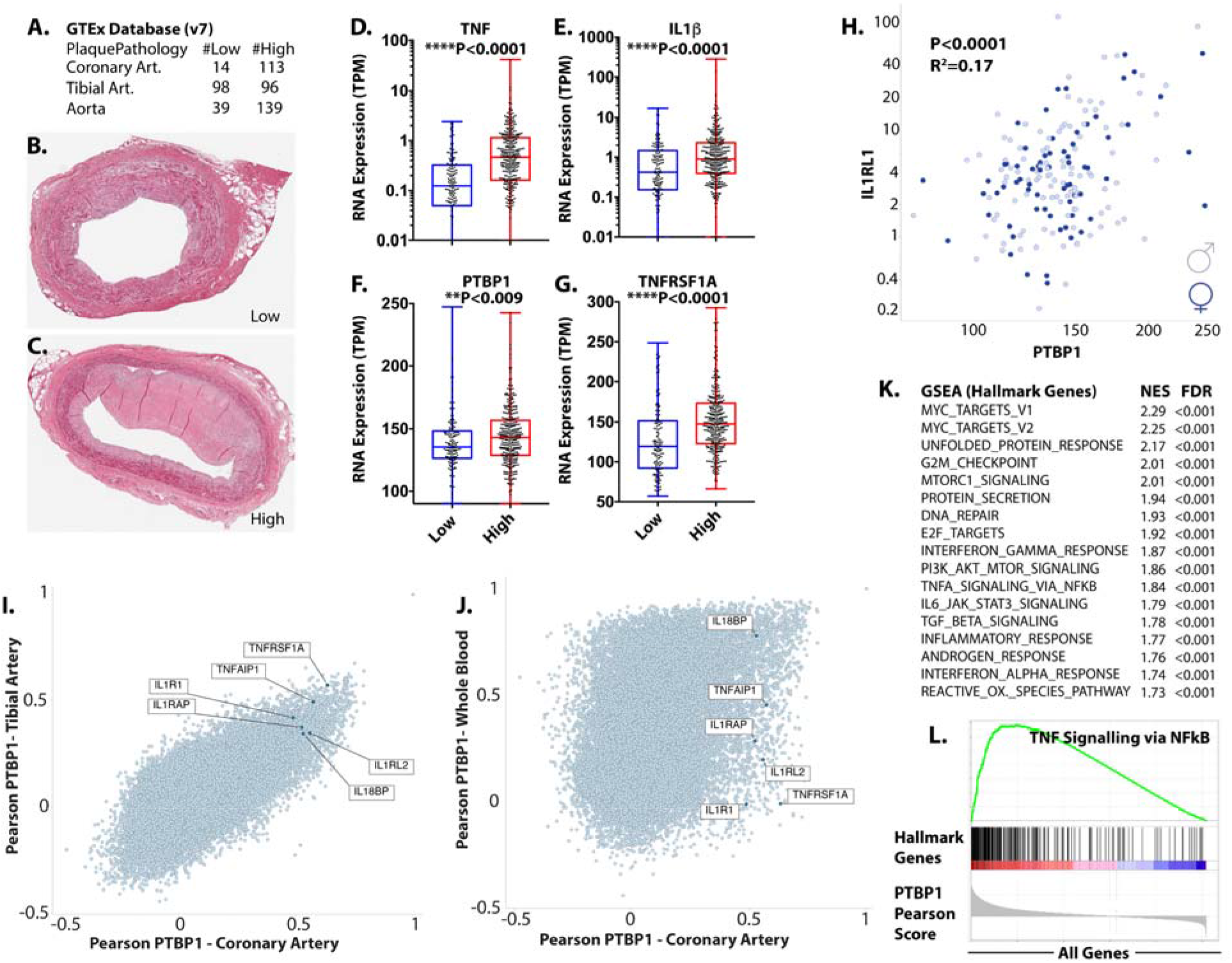
Ptbp1 expression correlates with plaque and expression of TNF pathway genes in human arteries. (A) Summary of arteries (Art.) of each type with indicated pathology data from the GTEx database (v7). (B&C) Examples of histological sections used for analysis of plaque characteristics in the GTEx database, showing examples of low (B) and high (C) plaque. (D-G) Analysis of expression of the indicated genes in arteries separated by low and high plaque phenotype. (H) Gene-gene correlation between the expression of PTBP1 and TNFRSF1A in human coronary arteries, where sex is indicated by different shading (gray=female; male=black). (I&J) Pearson correlation scoring for each gene-gene interaction in coronary and tibial arteries (I) or coronary arteries and blood (J), showing whether these gene-gene interactions are conserved between tissues. (K) Top GSEA Hallmark Gene Groups for genes most associated with Ptbp1 (highest Pearson scores). NES=normalized enrichment score, FDR=false-discovery rate (L) Example GSEA plot, showing the location of TNF Signaling via NFκB Hallmark Genes among those most associated with PTBP1 expression (left side of the Pearson Score plot). (D-G) Mann-Whitney test.

Thus, in human arteries, Ptbp1 expression is positively associated with plaque accumulation and with inflammatory pathways (TNF signaling via NFκB), that are also regulated by Ptbp1 in mice.

## Conclusions

Here, through a focused CRISPR-KO screen of splice factors involved in the early endothelial response to LDF, we found that Ptbp1 is required for efficient NFκB signaling responses in arterial endothelial cells. Absence of Ptbp1 limits NFκB transcriptional responses to pro-inflammatory cytokines by reducing nuclear translocation of NFκB, effects which are rescued by human PTBP1. As Ptbp1 expression and downstream splicing responses are induced in the arterial endothelium in the early inflammatory response to LDF, we propose that activation of Ptbp1 may be an important step in the priming of endothelial cells, licensing them for the induction of cell adhesion molecules Icam and Vcam upon cytokine stimulation. Consistent with this idea, Ptbp1 is positively correlated with human plaque burden and TNFα-mediated signaling pathways in human coronary arteries. Furthermore, deletion of Ptbp1 from the endothelium limits myeloid cell recruitment to sites of natural LDF and endothelial priming.

### Contextual role of Ptbp1 in priming acute and chronic inflammatory responses

In this work, we have used a simplified *in vitro* model to understand the role of RNA-binding protein expression on endothelial cell priming, and to show that this process depends on platelet recruitment. We have previously shown that endothelial cells cultured *in vitro* under serum and static conditions exhibit an expression and splicing profile that mimics activated carotid endothelium *in vivo*, after exposure to partial carotid ligation ^9^. This is an expected result, as typical *in vitro* culture conditions expose endothelial cells to platelet and immune releasate in serum, and a lack of flow that removes quiescence signals from laminar flow. Indeed, cultured endothelial cells *in vitro* are primed for the expression of Icam1 and Vcam in response to TNFα or Il1β stimulation ^5^. Expression of Ptbp1 is elevated *in vitro*, as it is *in vivo* in endothelial cells activated by exposure of LDF. Notably, the expression of Ptbp1 *in vivo* is not altered by flow alone, as it did not change under LDF conditions when platelets were removed. Similarly, although basal levels of Ptbp1 in culture were higher than *in vivo*, these levels were further increased by the addition of platelets and monocytes (SI Figure 7), but neither plasma nor platelets and monocytes alone. Thus, our model is that platelets are recruited to endothelial cells exposed to LDF *in vivo*, where they interact with the endothelium through an unknown mechanism to induce endothelial cell priming - mediated in part by an increase in Ptbp1 expression. This leads to an elevated response to pro-inflammatory cytokines, such as TNFα or Il1β, and increased recruitment of myeloid cells to the endothelium.

Our model is consistent with the observation that not all endothelial cells in LDF regions of aorta exhibit Vcam expression upon systemic LPS treatment ^4^. We propose that the differential response among cells may be due to local recruitment of platelets and activation of the priming response we describe here, through increased expression of Ptbp1. Consistent with this idea, platelets are critical for the recruitment of myeloid cells to regions of the vasculature exposed to LDF, and also for the development of atherosclerotic plaque^11–14^. While some of these effects are due to mechanisms previously described, such as deposition of CCL5 on the endothelial cell surface by activated platelets, and the tethering of myeloid cells to the endothelium by P-selectin expression on platelet intermediates ^14^ we propose that induction of Ptbp1 in the endothelium is an important consequence of platelet recruitment and contributes to the progression of vascular inflammation and plaque development.

### Ptbp1 and the NFκB signaling pathway

Polypyrimidine tract binding protein (Ptbp1, or hnRNP I) is widely expressed, and has been shown to regulate splicing responses and differentiation in a wide range of tissues, including neurons, cardiomyocytes and leukocytes. While this work is the first that we know of to show a direct effect on the NFκB signaling pathway, prior reports suggest that Ptbp1 effects on inflammatory responses are not limited to the endothelium. Ptbp1 was identified in an RNA interference (RNAi) screen to identify regulators of the senescence-associated secretory phenotype (SASP) in cancer cells, indicated by expression of IL-6 and IL-8 ^27^. NFκB signaling activation is critical in the induction of the SASP response, and small interfering RNA (siRNA) to NFκB subunit RELA suppressed the response as well. Interestingly, knockdown of Ptbp1 had little effect on the growth of cells in this screen, as we observed in human arterial endothelial cells here. However, in contrast to our results in endothelial cells, knockdown of Ptbp1 in cancer cells did not interfere with TNF-mediated induction of an NFκB reporter^27^. In another study, knockdown of Ptbp1 in T cells led to increased T cell expansion, which coincided with increased level of IKβα. While nuclear NFκB was not measured, the prediction would be that increased IKβα could lead to increased cytoplasmic sequestration of NFκB ^28^. The consistent theme is not the mechanism, as there was little difference in IKβα with or without Ptbp1 in arterial endothelial cells, but that the pathway is consistently affected by Ptbp1.

The specific mediators of the effect of Ptbp1 on the NFκB signaling pathway are not yet clear, but alterations in RNA splicing are a likely cause. RNA-binding proteins can have diverse cellular functions, including RNA transport, stability, and translation, but given the predominantly nuclear localization of the splice factor in these cells, altered nuclear splicing functions is the most likely possibility. Consistent with this idea, there is a substantial overlap between RNA splicing events regulated *in vivo* by the loss of platelets and those regulated *in vitro* by the loss of Ptbp1. Some notable examples include an altered 3’UTR in Iκβκγ (Nemo), a core NFκB signaling pathway component, a skipped exon in Pdlim7, an ubiquitin ligase of p65 which limits NFκB signaling^29^, and several genes linked to autoimmune or inflammatory diseases in human genetic association studies, such as STAT3 (Crohn’s disease, multiple sclerosis, and psoriasis)^30^, FNBP1 (Crohn’s disease, multiple sclerosis, psoriasis, Type I diabetes)^31^ and MAGI1 (Crohn’s disease, psoriasis and Type-1 diabetes)^32^. In total, we detected splicing alterations in about half of the Kegg NFκB genes in our Ptbp1 KO cells, and many more splicing events peripherally related to the pathway. It is likely that the reduced NFκB signaling response we observed in cells is a result of more than one of these alterations, but a future broad analysis of the regulated splicing events will be required to understand their relative contributions.

### Ptbp1 in the context of atherosclerosis and human cardiovascular disease

Two major risk factors for atherosclerosis are also linked to increased Ptbp1 expression: senescence and integrin-mediated adhesions to fibronectin. Increased Ptbp1 expression has been observed with age and in senescent cells ^27,33^. Aging is arguably the single greatest risk factor for atherosclerosis. Increasing numbers of senescent cells in the arterial intima appear to be an important contributor^34^. Senescent endothelial cells express the potent proinflammatory cytokine IL6 as a result of increased NFκB activity^35^, and markers of senescence are more abundant in patients with endothelial dysfunction ^36^. Ptbp1 localization is also dynamically altered by cell adhesion to fibronectin matrix ^37^, being mainly in the cytoplasm during adhesion and until the cell has established focal adhesions – at which point its nuclear localization is restored. In mouse embryonic fibroblasts, Ptbp1 was bound to mRNA encoding vinculin and alpha-actinin – bringing them to the spreading periphery, and was required for cell spreading ^37^. In response to LDF patterns and in early atherogenesis, fibronectin is deposited beneath the endothelium of arteries^6,10,38^. Fibronectin that is deposited, either from the plasma or the endothelium, promotes NFκB activity and myeloid recruitment and plaque in these regions ^6,39^. As these responses are determined by the specific matrix binding integrins ^6,40^, it may be interesting to examine Ptbp1 localization and activity upon binding of cells to other matrices than fibronectin.

Our evidence from cells and the animal model indicates a causal role for Ptbp1 in endothelial activation and plaque development. While there have been no focused studies on Ptbp1 function in human atherosclerosis or inflammatory diseases, we note that a SNP in Ptbp1 is correlated with C-reactive protein (CRP) levels, a marker of systemic inflammatory responses (rs123698-G, P=1e-9) ^41^. As Ptbp1 levels in human arteries correlated with more severe plaque, and perhaps more importantly, with RNA markers of TNFα-NFκB signaling pathway activity in those plaques, understanding the links between innate immune cell recruitment, senescence, and matrix adhesion, and Ptbp1 activity may provide new insights and approaches to understanding inflammatory risk in atherosclerosis and other inflammatory diseases.

In conclusion, we report that Ptbp1 – with increased in expression in regions of the vasculature exposed to low and disturbed flow – is required for NFκB nuclear localization upon stimulation. As loss of Ptbp1 reverts alternative splicing patterns in cultured and activated endothelial cells towards a quiescent endothelium, and blunts the subsequent response to cytokine stimulations by TNFα and Il1β, we propose that Ptbp1 is a critical component of endothelial cell priming. We predict that this Ptbp1 function will be conserved across different vascular beds and inflammatory responses. Therefore, further investigation of this mechanism leading to increased expression and activity of Ptbp1, its associated RNA-binding splice factors, and the RNA transcripts they regulate is warranted.

## Materials & Methods

### CRISPR-KO Screening and Single Cell Clones

#### Generation and amplification of CRISPR plasmid pools

(Splice factor pool) Five CRISPR-KO guide sequences were designed to each of 57 splice factor targets using described rules (Azimuth 2.0) ^42^. Oligos were ordered and BsmBI batch-cloned into the gRNA expression core of a LentiCRISPR_v2 plasmid containing Cas9 (Addgene 52961). The library was amplified in Stbl3 (Thermo) and tested by miSeq analysis for coverage and skew. Individual guides (e.g. to Rbfox2) followed the same protocol, but were confirmed by Sanger sequencing. (Genome-wide pool) For the genome-wide screen, the Brie library in LentiCRISPR_v2 (Addgene 73632) was obtained as a plasmid library and amplified in Stbl4 (Thermo) on 500cm^2^ LB-agar coated ampicillin+ selection plates. The library was tested by NextSeq analysis for coverage and skew.

#### Transduction of Splice Factor pool and Genome-Wide CRISPR pool into mouse aortic endothelial cells (mAEC)

Lentivirus was generated in 293T cells using delta 8.9 and pHCMV-EcoEnv (Addgene 15802) as packaging plasmids and 25kDa linear PEI as a transfection agent. Media was changed at 1 day and supernatant was taken at 3 days after transfection for the treatment of recipient cells at MOI <0.3 (>1000 infected cells per guide). Virus supernatant was added to recipient mAECs with polybrene (8ug/mL), and cells were selected with puromycin for 4 days, confirming a MOI<0.3 and complete killing of uninfected cells. In experiments examining induction of Icam and Vcam in the endothelium and protein and RNA analysis, recipient cells were TetOn-Sv40 aortic endothelial cells, prepared as previously described ^9,10^. In experiments in which eGFP reported NFkB signaling activity, recipient cells had been infected with a modified version of an NFkB reporter construct, engineered to express RFP constitutively and eGFP upon activation of the 4x NFkB binding motif. An aliquot of cells was taken 3-4 days post infection for an early timepoint analysis of lentiviral library representation.

#### Assessing effects on inflammatory response

mAECs harboring CRISPR guides were trypsinized and split into low glucose DMEM and 10% FBS into new dishes, without doxycycline to stop growth. 24 hours later, cells were once again trypsinized and collected for staining and flow cytometry.

### Enrichment of guides in cells with increased or decreased Icam1/Vcam or NFκB reporter

DNA was collected from sorted subsets of cells Quick DNA kit (Zymo). Lentiviral insertions in genomic DNA were amplified by nested PCR, using PCR1 primers (see table) followed by PCR2 primers containing barcodes and Illumina priming sequences (see table). PCR reactions were separated using a 2% agarose gel, and bands of interest were excised. PCR products were column cleaned (Gel cleanup kit, Zymo), and pooled based on relative concentration. Pooled samples were sequenced on Illumina NextSeq instrument using 75bp single end (SE) reads.

### Identification of top Icam1/Vcam or NFκB regulators (Bioinformatics approach)

Fastq files were analyzed using Mageck-0.5.6 software, providing counts of each guide in each sequenced population, and differences in their representation (mageck test).

### Expression of human PTBP1 cDNA in cells

Poly(A) primed cDNA was prepared from human aortic endothelial cells. PCR primers with restriction sites were used to amplify the expected band size for Ptbp1, and this was ligated into pLV-EF1a-IRES-Blast (Addgene Plasmid #85133) using EcoR1 and BamHI restriction sites. The sequence of the inserted cDNA was confirmed by Sanger sequencing. In the described experiments, both a “empty” rescue construct and the Ptbp1 containing construct were used.

### Immunofluorescence and Quantification

mAEC (TetOn-SV40, hTert immortalized) were plated onto collagen-coated coverslips in a 6 well cell culture plate without the presence of doxycycline (to stop growth), but with low glucose DMEM + 10% FBS. 24 hours after plating, DMEM + FBS was removed, and coverslips were washed in the cell culture plates with cold PBS (3x). Cells were fixed with cold 4% paraformaldehyde (PFA) for 5 minutes on ice. Coverslips were washed again 3x with cold PBS. For Ptbp1 staining, cells on coverslips were permeabilized for 10 minutes at room temperature with 0.1% Triton-x in PBS. All other immunofluorescence experiments did not permeabilize before staining with primary antibody. Blocking buffer for Ptbp1 staining was 1% BSA, 0.3M glycine in PBS + 0.1% Tween for 1 hour at room temperature. All other staining used 10% normal goat serum (NGS) + 0.1% Tween in PBS. Ptbp1 staining was performed for 2 hours at 1:500 (Abcam, ab133734) and NFκB (Cell Signaling, 8242) staining for 2 hours at 1:100 at room temperature. Secondary staining was done for 1 hour at room temperature using a fluorescently-conjugated antibody. Images were taken on a Zeiss epifluorescent microscope using ZenPro software. Quantification was performed using ImageJ (1.50i).

### Western Blot and Quantitation

2x Laemmli buffer was added to the adherent cells and 2uL of phosphatase inhibitor cocktail 2 (Sigma, P5726) and 2uL of phosphatase inhibitor cocktail 3 (Sigma, P0044) were added to the cell culture plates for 5 minutes before cell lysate was scraped into 1.5mL Eppendorf tubes. 1uL of Benzonase (Sigma, E1014) was added to each sample and lysates were placed on ice for 5 minutes before being sheared 5 times with a 26 gauge needle. 5% of total volume was then placed in a separate Eppendorf tube, and freshly prepared 1M DTT (dithiothreitol, 1uL/5uL cell lysate, Sigma D9779) was added. Samples were then placed at 95°C for 5-10 minutes before being loaded onto the western blot gel. Gels were run at 125v for 1.5 hours before western blot transfer was set up to run overnight at 4°C (100v for 1 hour, then O/N at 30v). Blocking was done using 5% milk in TBST. Primary antibody staining (dilution is antibody dependent, but followed manufacturer’s recommendation) was done for 2 hours at room temperature, secondary staining (1:5,000) was done for 1 hour at room temperature, and GAPDH staining (1:10,000) was done for 2 hours at room temperature, all with rotation. Blots were imaged using Clarity ECL Western Blot substrate (Biorad, 1705060). Quantitation was performed using ImageJ (1.50i) by inverting the Western Blot image and subtracting the background intensity from the GAPDH and the band of interest intensity.

### Nuclear Fractionation

Adherent cells were trypsinized and pelleted. Cell counts were performed prior to any further experimentation. Each cell pellet was resuspended in 100uL of Nuclei EZ Prep Isolation buffer (Sigma, NUC101) and placed on ice for 5 minutes before being centrifuged (5 minutes at 500 x G). Supernatant (cytoplasmic fraction) was collected, and to nuclei fraction, another 500uL of EZ Prep Isolation buffer was added (resuspending the cell pellet in the buffer) and placed on ice for 5 minutes before 500uL of Isolation buffer was added (for 1 mL total volume). Cells were pelleted by centrifugation, supernatant was removed, and pellet was resuspended in 100uL of EZ Prep Isolation buffer before being prepared for western blot analysis.

### Mice

For the endothelial deletion of Ptbp1, *Cdh5(PAC)-CreERT2* and *Ptbp1^lox/lox^* mice were used, which have been previously described ^2221^. They were intercrossed to create the Ptbp1 EC-KO mice (*Cdh5(PAC)-CreERT2; Ptbp1^lox/lox^*)and littermate controls (*Ptbp1^lox/lox^*) used here. Mice were used between 2 and 7 months of age in paired groups of males and females. Tamoxifen (Sigma) was delivered intraperitoneally, dissolved at 10mg/mL in sunflower oil. 1mg was given in each of three doses.

All mice were housed and handled in accordance with protocols approved by the University of Connecticut Health Center for Comparative Medicine.

### Hyperlipidemia

Mice received 100 μl intraperitoneal injections of 1 × 10^11^ viral particles of AAV8-encoding mutant PCSK9 (pAAV/D377Y-mPCSK9) produced at the Gene Transfer Vector Core (Grousbeck Gene Therapy Center, Harvard Medical School). Mice were then placed on the Clinton/Cybulsky high-fat rodent diet (HFD) with regular casein and 1.25% added cholesterol (D12108C; Research Diets). To measure cholesterol levels, blood was collected from the right ventricle into lithium-heparinized tubes (365965; BD Biosciences) and centrifuged at 5,000 *g* for 10 min to obtain serum. Samples were stored at −80°C and analyzed by the Total Cholesterol Assay Kit (INC, cat. no. STA-384; Cell Biolabs).

### Aortic digestion

Vessels were flushed with phosphate-buffered saline (PBS) through the left ventricle and out the right atrium and then dissected free of adventitial tissue, minced with scissors into 1.5 ml microcentrifuge tubes (Eppendorf), and incubated for 1 hour at 37°C with gentle rotation (20 rpm) in balanced salt solution (BSS) media containing 150 U/ml collagenase type IV (Sigma-Aldrich C5138), 60 U/ml DNase I (Sigma-Aldrich), 1 μM MgCl_2_ (Sigma Aldrich), 1 μM CaCl_2_ (Sigma-Aldrich), and 5% fetal bovine serum (FBS). Digested tissues were crushed through 35 μm cell-strainer caps (BD Biosciences) and quenched with 5 ml of cold BSS + 10% FBS in round-bottom tubes. Supernatant was removed after a 5 minute, 320 *g* centrifugation, and the cell pellet was resuspended and quantified using a Z1 particle counter (Beckman Coulter).

### Flow cytometry

#### Aortic digestion and analysis

Vessels were flushed with phosphate-buffered saline (PBS) through the left ventricle and out the right atrium and then dissected free of adventitial tissue, minced with scissors into 1.5-ml Eppendorfs, and incubated for 1 h at 37°C with gentle rotation (20 rpm) in balanced salt solution (BSS) media containing 150 U/ml collagenase type IV (Sigma-Aldrich C5138), 60 U/ml DNase I (Sigma-Aldrich), 1 μM MgCl_2_ (Sigma Aldrich), 1 μM CaCl_2_ (Sigma-Aldrich), and 5% fetal bovine serum (FBS). Digested tissues were crushed through 35-μm cell-strainer caps (BD Biosciences) and quenched with 5 ml of cold BSS + 10% FBS in round-bottom tubes. Supernatant was removed after a 5-min, 320-g centrifuge, and the cell pellet was resuspended and quantified using a Z1 particle counter (Beckman Coulter).

#### Flow Cytometry Staining

Samples were stained in 2%FBS with 1mM EDTA in PBS, with the following: LIVE/DEAD UV Blue (1:100, L34962, ThermoFisher), CD8 (1:200, Biolegend, 100708, clone 53-6.7), CD4 (1:200, Biolegend 100536, clone RM4-5), CD45.2 (1:200, Biolegend, 103116, clone 30-F1111), Cd3e (1:200, Biolegend, 100348, clone 145-2C11), Gr1 (1:200, B.D. Pharmingen, 552093, clone RB6-8C5), and Cd11b (1:200, Biolegend, 10112, clone M1/70). LSR Aria-IIA (BD Biosciences) was used for acquisition. Viable cell gate is representative of a size gate, single-cell gate, and viability gate.

Acquisition was performed on an LSR analyzer (Becton Dickinson). All flow cytometry data were analyzed with FlowJo (Tree Star, Ashland, OR).

#### Histology

Innominate arteries were harvested and placed cores before being placed in Zinc-formalin fix for 24 hours. 24 hours later, cores were moved into 70% ethanol solution. Innominate arteries were then paraffin infused, before being embedded in paraffin. 5um tissue sections were cut for each mouse. Lung tissues were harvested and placed in PBS before being put in 4% paraformaldehyde (PFA) + 5% sucrose overnight at 4°C. Tissues were then embedded in OCT before being sectioned in 5um sections.

#### Analysis of GTEx data

GTEx (version 7) was used ^43^. Pathology was determined from the provided sample tables. Terms “some intimal thickening”, “no lesions”, “no plaques”, “no significant atherosclerosis” were linked to the group “Low Plaque”. Terms “mild plaques”, “minimal plaques”, “mild atherosclerosis”, “atherosclerosis”, “atherosis”, etc. were linked to the group “High Plaque”. Gene-gene correlations were performed in R using gene level transcript expression table, using the “cor” function. Gene set enrichment analysis was performed using the desktop version of GSEA and a preranked list of genes correlated with Ptbp1 (most to least), as a weighted analysis ^44^.

#### Analysis of Gene Expression and Splicing

RNA was isolated from fixed nuclei using an RNAeasy kit (RNAeasy, Qiagen 74104) with on column DNAse treatment. For RNA-sequencing, samples were prepared for library prepraration using TruSeq RNA Library Prep Kit v2 (Illumina). Total RNA was quantified and purity ratios determined for each sample using the NanoDrop 2000 spectrophotometer (Thermo Fisher Scientific, Waltham, MA, USA). To further assess RNA quality, total RNA was analyzed on the Agilent TapeStation 4200 (Agilent Technologies, Santa Clara, CA, USA) using the RNA High Sensitivity assay. Amplified libraries were validated for length and adapter dimer removal using the Agilent TapeStation 4200 D1000 High Sensitivity assay (Agilent Technologies, Santa Clara, CA, USA) then quantified and normalized using the dsDNA High Sensitivity Assay for Qubit 3.0 (Life Technologies, Carlsbad, CA, USA).

Sample libraries were prepared for Illumina sequencing by denaturing and diluting the libraries per manufacturer’s protocol (Illumina, San Diego, CA, USA). All samples were pooled into one sequencing pool, equally normalized, and run as one sample pool across the Illumina NovaSeq. Target read depth was achieved per sample with paired end 150bp reads.

Paired-end FASTQ files were processed using Whippet (Julia 0.6.4 and Whippet v0.11) using default settings, after generating an index from GRCm38.primary_assembly.genome.fa.gz and gencode.vM23.annotation.gtf.gz (Gencode, M23). Psi files and read support were used to select only splicing events with read coverage >5 across all replicates. TpM files were used to examine expression level. ggSashimi was used to plot alternative splicing events identified by Whippet analysis.

## Key Resources Table

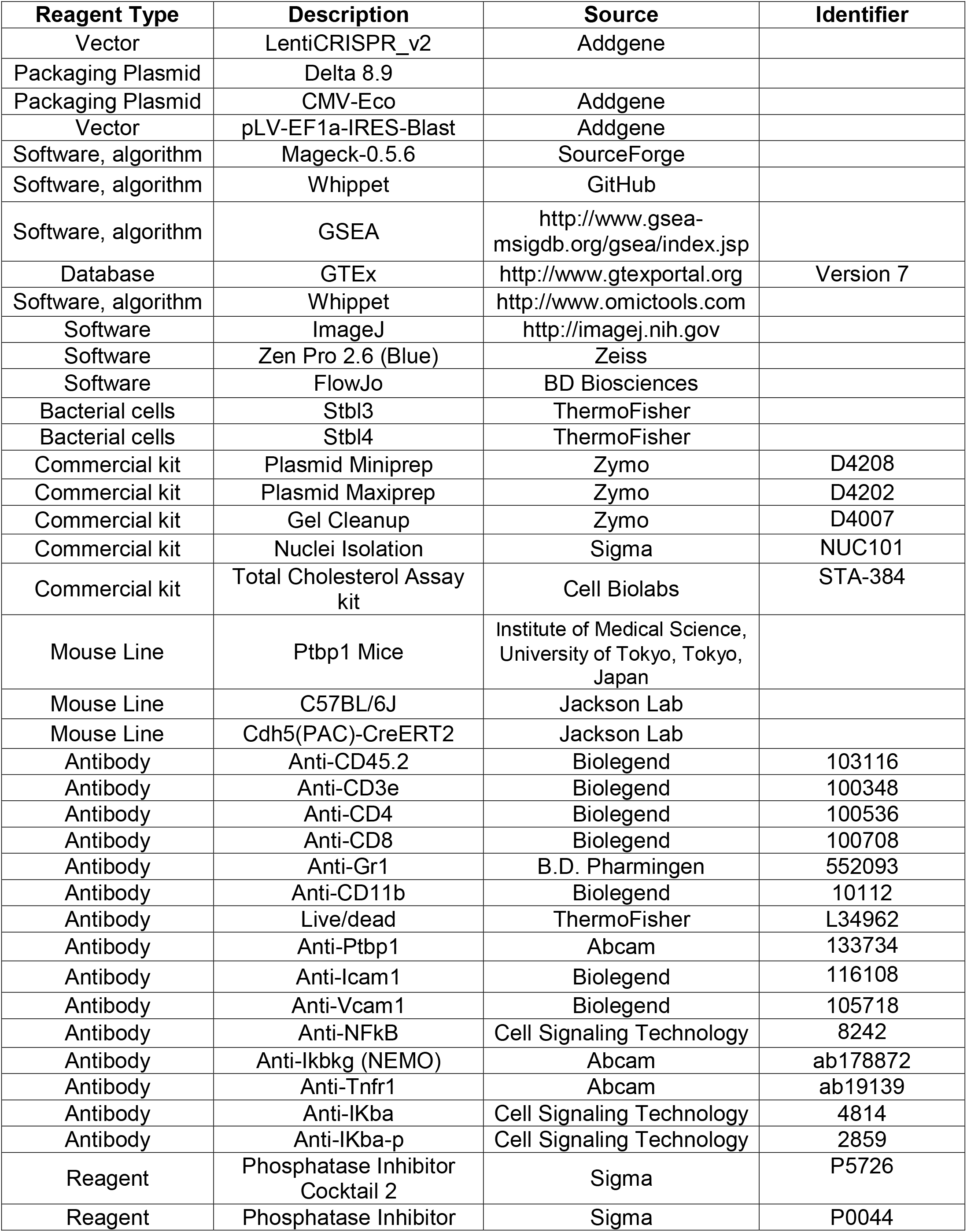
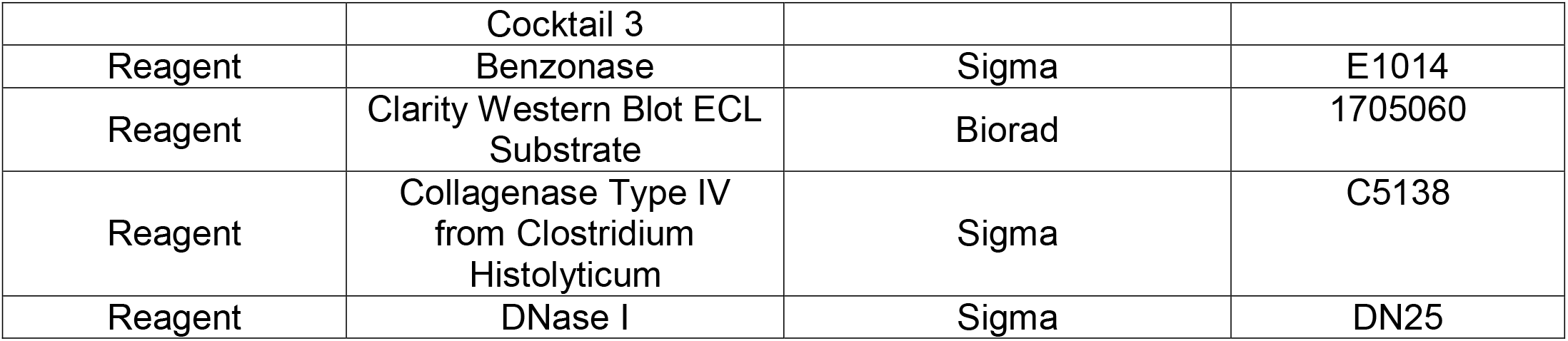

## Acknowledgements

We appreciate the work of Xenia Bradley, who worked with Jessica in the summer of 2017 to test conditions for the CRISPR screen that was eventually performed. Bo Reese, in the Center for Genome Innovation at UCONN aided in the adaptation of the CRISPR screen sequencing protocols from published work, and was a valuable consult in RNA-sequencing studies performed here. In the course of this work, we received valuable input from the ImmunoCardiovascular Group at UCONN Health, particularly Dr. Beiyan Zhou (University of Connecticut) and Alison Kohan (University of Pittsburgh). We also thank Christopher Bonin and Geneva Hargis in the Science Writing and Illustration group at UCONN Health, for editing and help with our graphical abstract.

