## Supplemental Figures for "Splice Factor Polypyrimidine tract-binding protein 1 (Ptbp1) is Required for Immune Priming of the Endothelium in Atherogenic Disturbed Flow Conditions"

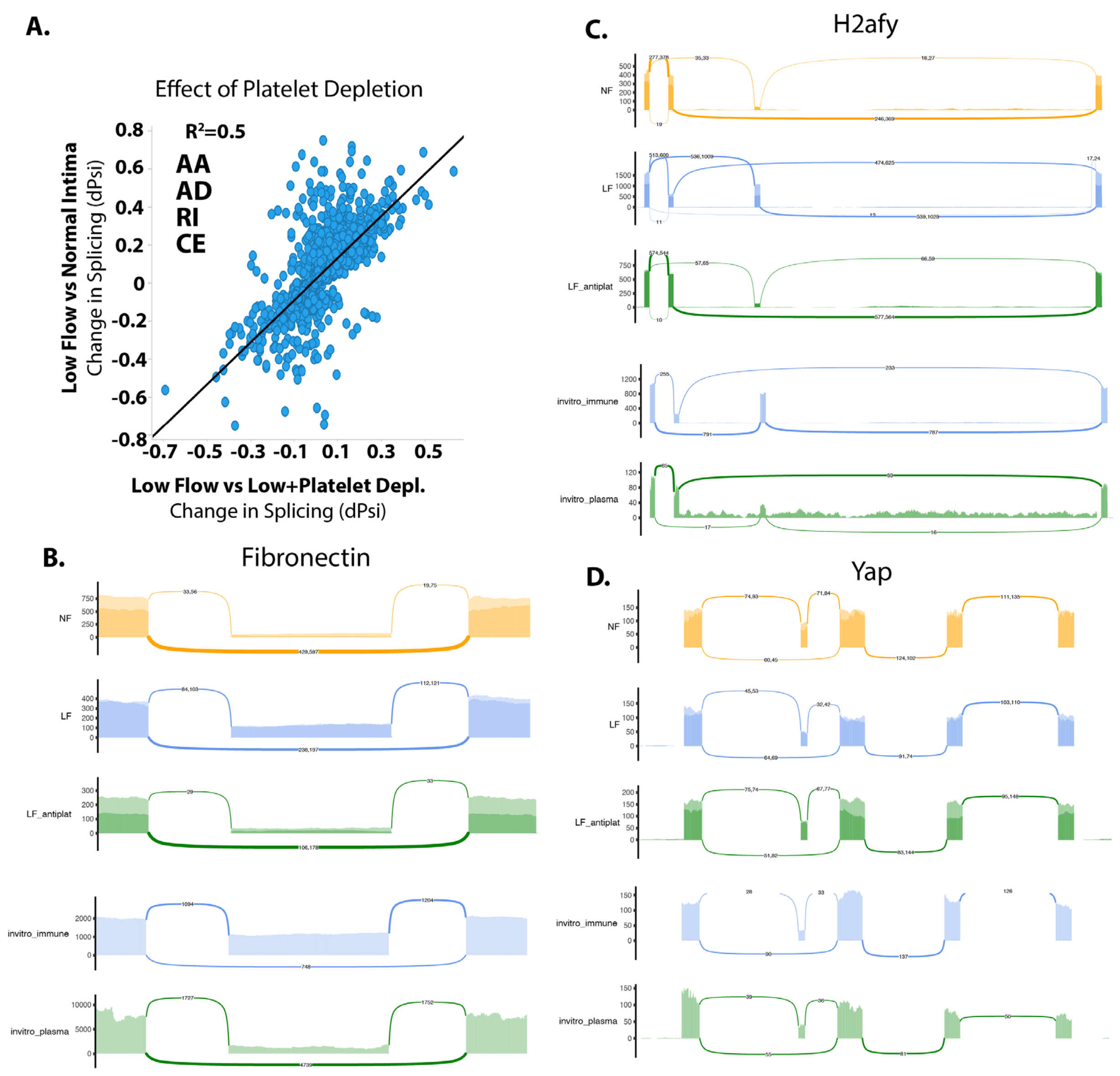


**SI Figure 1. Platelet depletion reverses splicing changes induced by low and disturbed flow (LDF).**
(A) Graph showing the change in inclusion level (Psi) for the indicated classes of splicing events, in a comparison of LDF versus normal flow intima (y axis) or low flow intima versus low flow intima with platelet depletion (x axis). (B-D) Sashimi plots showing example splicing changes altered under LDF conditions (LF) relative to normal flow conditions (NF) and reverted under LDF conditions after the depletion of platelets (LP_antiplat). In addition, the plots show the effect of adding a combination of platelets and monocytes to endothelial cells in vitro (invitro_immune), versus plasma alone (invitro_plasma).

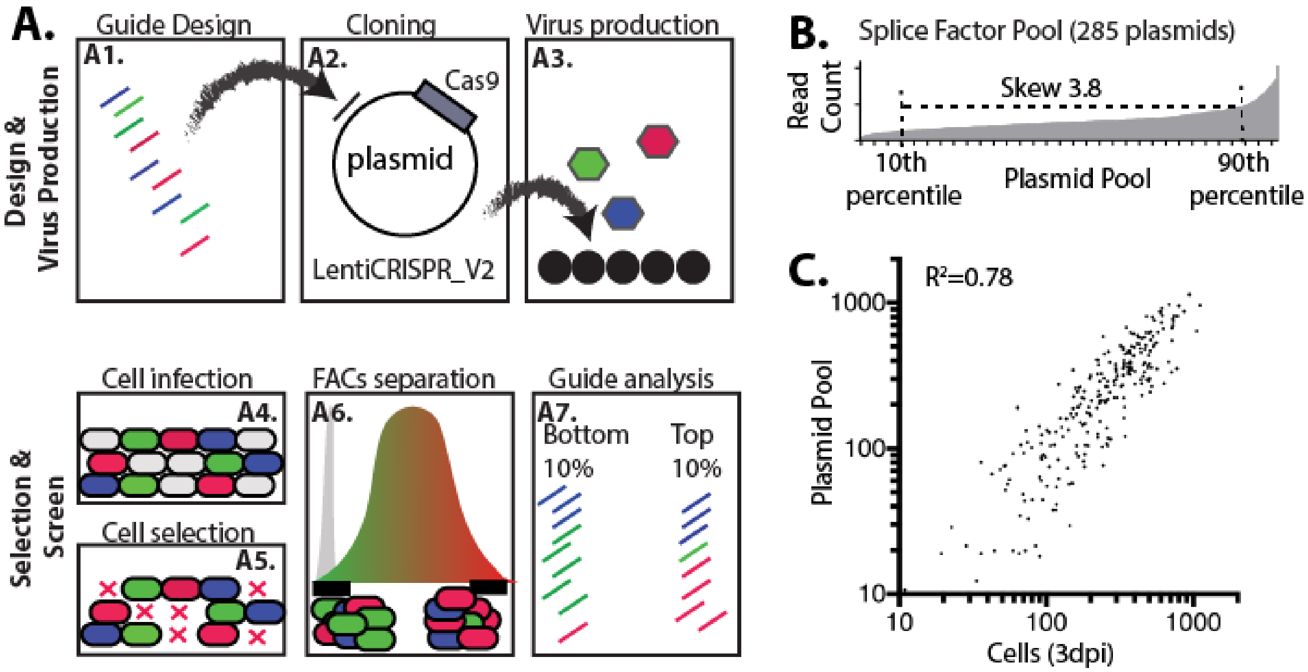


**SI Figure 2. Showing CRISPR KO screening approach.**

(A) Screening approach (A1) Guides to each target gene of interest were designed using the latest rules in the Broad design tool, and as hybridized oligos for (A2) batch cloning into a lentiviral backbone containing Cas9 and guide expression site. (A3) The resultant plasmid library is then used for production of lentivirus and infection of cells (A4). Infection is at 0.3 multiplicity of infection (MOI) and (A5) selection is by puromycin. (A6) Selected cells are sorted on the inclusion of alternative exon. Cells in the top 10% and bottom 10% of the response are isolated. (A7) Genomic DNA is isolated and guide enrichment in cells is assessed by sequencing of amplified lentiviral insertions. (B) Read density (y-axis) for each guide construct in the plasmid library for a list of 57 candidate splice factors (5 guides for each gene, x-axis). (C) Plasmid pool is generated as lentivirus and integrated into cells at similar levels. dpi=days post infection.

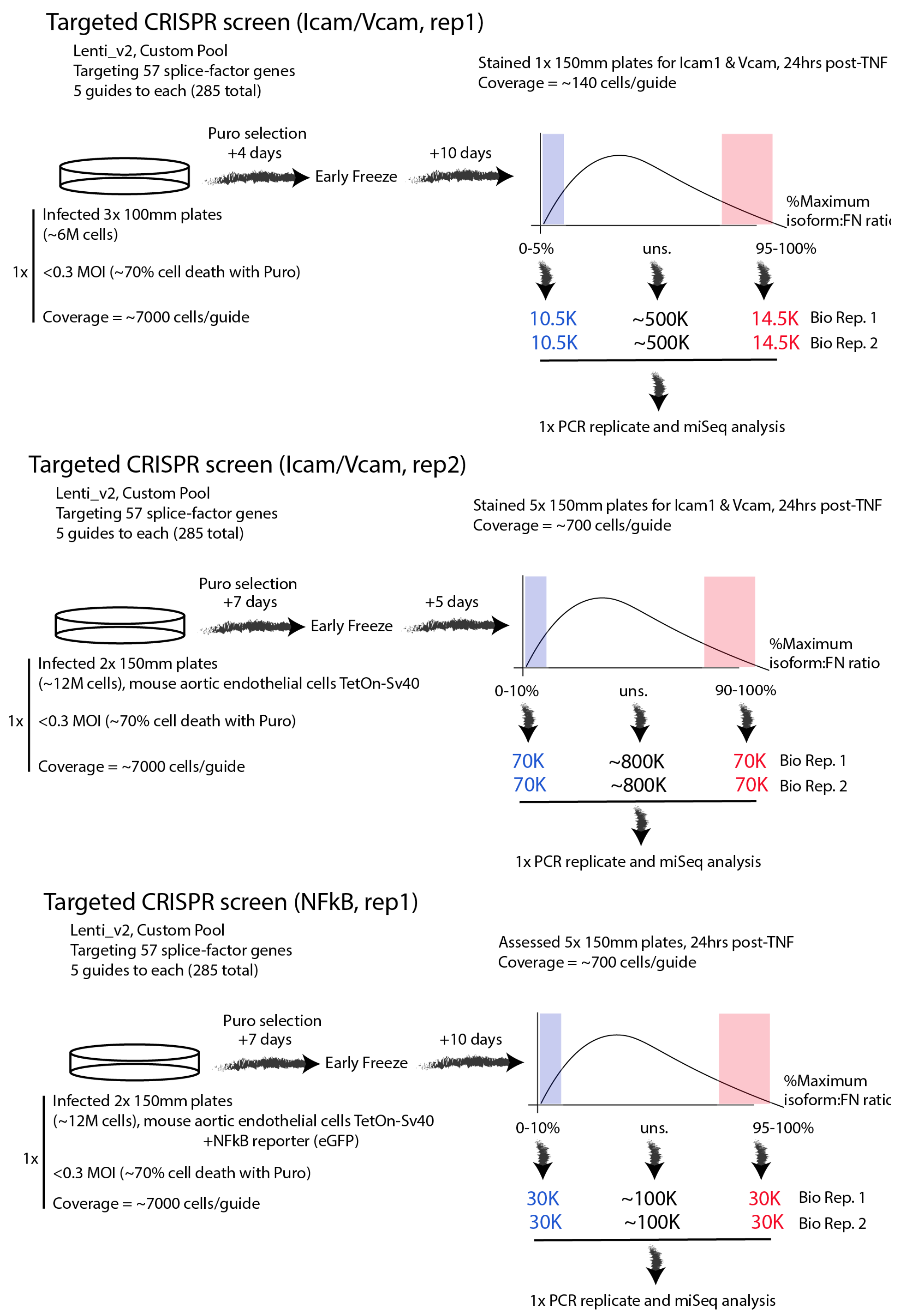


**SI Figure 3. CRISPR screen coverage.**
Schematic showing the preparation and analysis of cells in each individual CRISPR-KO screen performed. Numbers underneath the schematic of the flow-cytometry plot indicate the numbers of sorted cells from the high, low or unsorted fractions used in each biological replicate.

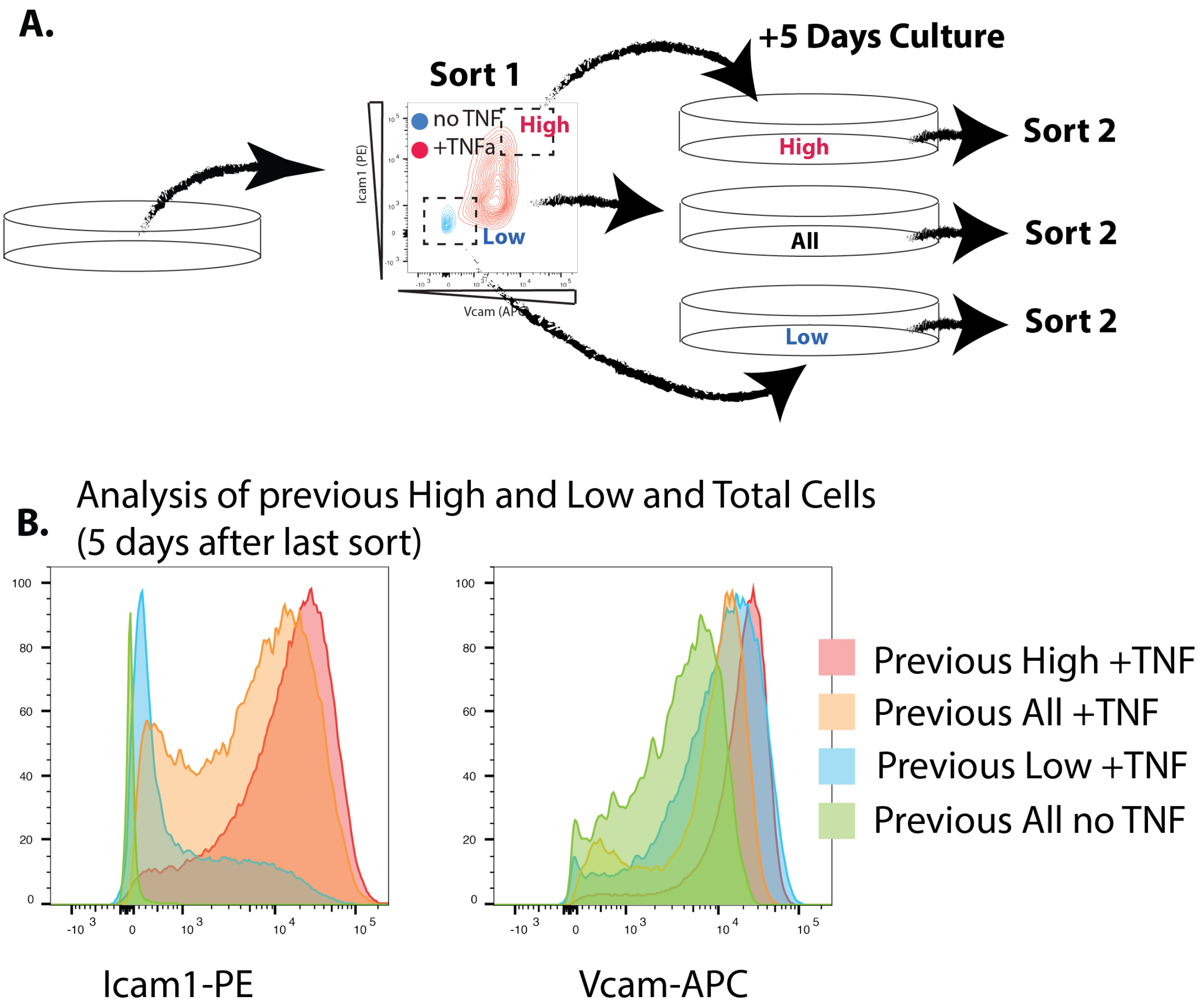


**SI Figure 4. Retention of Icam/Vcam response in sorted high and low responder populations.**
(A) Schematic showing secondary analysis sorted cells, to confirm the retention of low or high Icam/Vcam sorted populations in a second response to TNF$\alpha$ treatment. (B) Modal distribution of previously sorted Icam/Vcam high (High) and Icam/Vcam low (Low) populations, and the total population (All).

A.

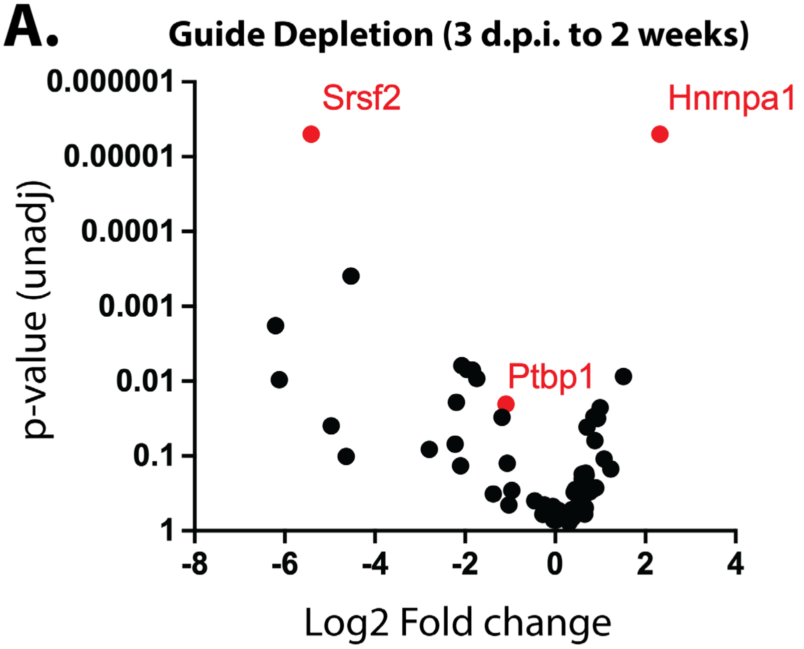


B.

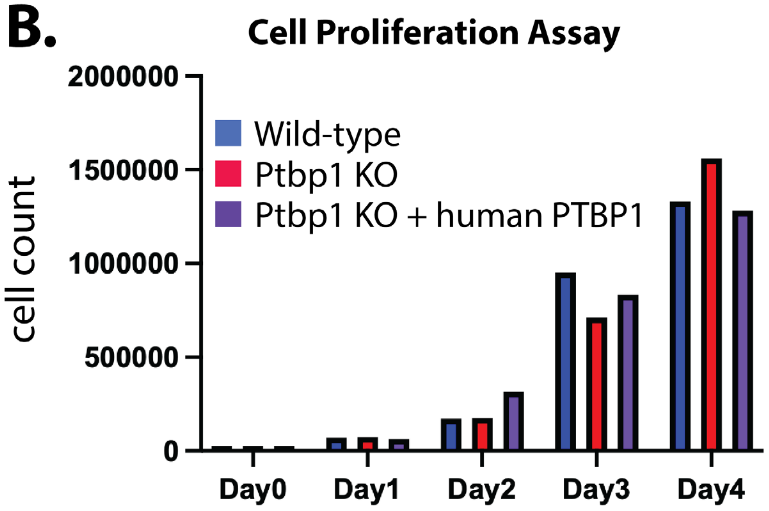


**SI Figure 5. Ptbp1 regulation of cell viability and proliferation.**
(A) Volcano plot showing guide depletion over two weeks of culture, from the initial infection of cells. d.p.i.=days post infection. Each gene is represented by 5 guides, which are pooled in this gene level analysis by MAGeCK. (B) Proliferation rates of wild-type aortic endothelial cells (TetOn-Sv40 in 2 μg/mL Dox media) or these cells with Ptbp1 CRISPR-KO, or rescue with human PTPB1 cDNA in the KO cells.

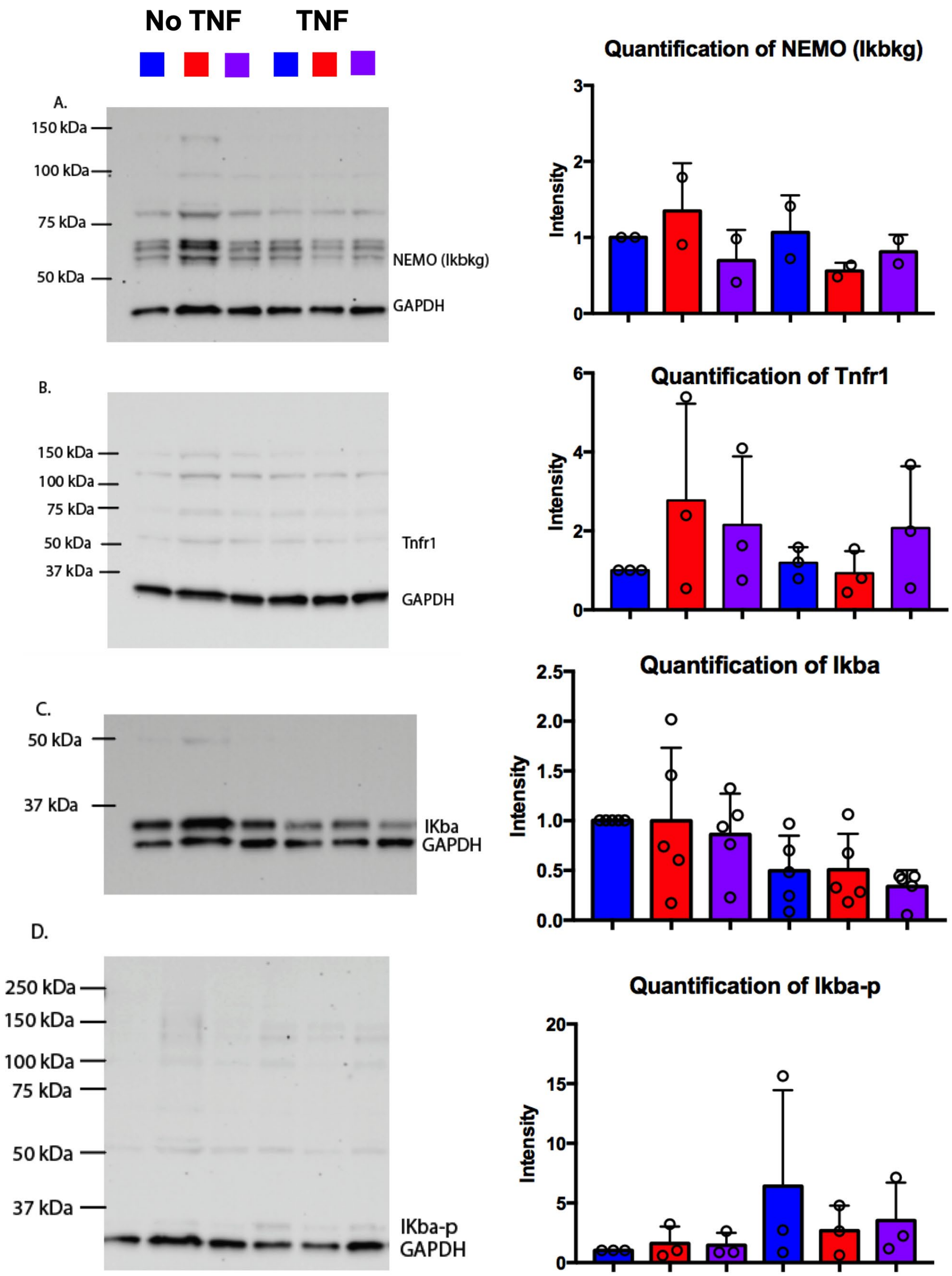


**SI Figure 6. Analysis of core NFκB pathway components.**
(A-D) Western blots and quantitation of signal intensities, relative to GAPDH control from biological replicates of the indicated cell lines (wild-type aortic endothelial cells (blue), Ptbp1 CRISPR-KO (red) and KO with human PTBP1 cDNA (purple), without (left) or with (right) 30ng/mL TNFα.

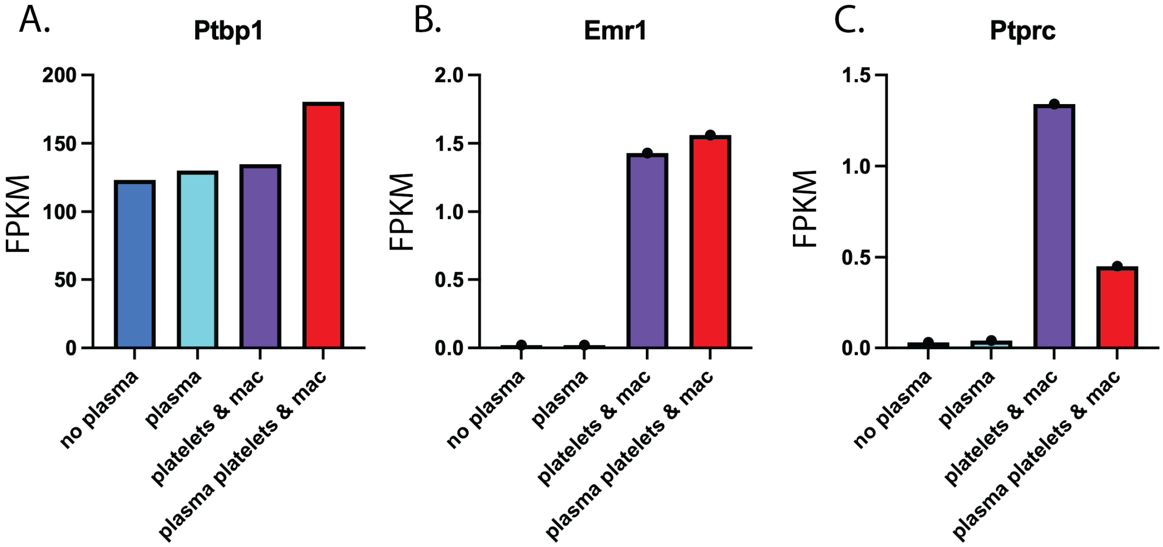


**SI Figure 7. Addition of innate immune cells and plasma increases Ptbp1 expression *in vitro*.**
Expression of the indicated genes in cultured aortic endothelial cells, in DMEM media only (dark blue), in DMEM with plasma (light blue), in DMEM with platelets and bone marrow derived monocytes/macrophages (mac, purple), or in DMEM with plasma and platelets and bone marrow derived monocytes/macrophages (red). Emr1 (F4/80) and Ptprc (CD45) are markers of macrophages and hematopoietic cells, respectively.
